# Expansion, functional diversification and gene fusion events in the Ato protein family

**DOI:** 10.1101/2025.05.26.656167

**Authors:** Faezeh Ghasemi, Cláudia Barata-Antunes, Yiannis Pyrris, Patrícia Ataíde, João Alves, Vitor Fernandes, Rosana Alves, Alexandra Gomes-Gonçalves, Margarida Casal, Wouter Van Genechten, Jana Nysten, Alistair J. P. Brown, Patrick Van Dijck, George Diallinas, Alexandros A Pittis, Isabel Soares-Silva, Sandra Paiva

## Abstract

*Candida albicans*, a commensal opportunistic pathogen, exhibits remarkable metabolic flexibility and adaptability to environmental changes. In glucose-limited niches, it utilizes alternative carbon sources such as carboxylic acids, which may influence its pathogenicity. In *Saccharomyces cerevisiae*, the uptake of monocarboxylates occurs through regulated plasma membrane (PM) transport proteins, such as Ato1 (Ady2), which belongs to the Acetate Uptake Transporter (AceTr) family. In *C. albicans*, these proteins are notably expanded, consisting of ten Ato-like proteins (ATO1-ATO10), whose functions remain unknown. Here, we investigated the role of Ato proteins in carboxylic acid utilization by *C. albicans* using *in-silico* and functional analysis. Our data revealed that several *C. albicans* Atos retain conserved AceTr motifs but possess distinct structural features, including differences in pore radius and binding sites for acetate and lactate. Expression analysis revealed that Ato1, Ato2, Ato3, and Ato6 exhibit distinct cellular localization and expression levels on the plasma membrane, depending on the presence or absence of monocarboxylates. Remarkably, deletion of *ATO1* impaired Ato2 and Ato3 expression and caused ER retention of a distinct form of Ato2, suggesting a central regulatory role for Ato1 in the Ato transport system. Finally, we identified a novel Ato-related protein family in vertebrates. This family has three consecutive 6-helix transport domains and a unique C-terminal fusion with Sua5/YciO/YrdC, an enzyme involved in tRNA modification.

Overall, our data suggests that the Ato protein family might play a critical role in the utilization of acetic or lactic acids in *C. albicans*. It also proposes potential functional redundancy among its members, which may contribute to rapid environmental adaptation and pathogenicity.

## 1. Introduction

Opportunistic fungal pathogens of humans include several *Candida* species that have become serious threats to global public health (1–4). These fungi can cause a broad spectrum of diseases, ranging from superficial infections of the vaginal and oral mucosa to life-threatening systemic infections (5,6). The genus *Candida* comprises 200 species, with approximately 10% identified as human pathogens (7). The most clinically significant species include *C. albicans*, *C. glabrata*, C. *auris*, *C. tropicalis*, *C. parapsilosis* and *C. krusei* (8–12). *C. albicans,* a commensal microorganism, exhibits the capacity to colonize various niches of the host, such as skin, oral cavity, gastrointestinal tract, and urogenital tract (13–15). Considerable metabolic flexibility underlies this adaptability (16,17).

Although sugars are preferred carbon sources for *C. albicans* (18), this pathogen can rapidly adapt to glucose-limited environments, such as the vaginal and gastrointestinal tracts (19,20). In these niches, *C. albicans* cells can utilize alternative carbon sources such as carboxylic acids (lactate, acetate, succinate, butyrate or propionate), amino acids and N-acetylglucosamine, which are produced by host cells or the resident microbiota (16,20–23). Carboxylic acids exist in two forms depending on their pKa and the ambient pH. When the environmental pH is lower than the pKa, the protonated (undissociated) form of acid can cross the plasma membrane by passive diffusion. Conversely, when the pH exceeds the acid’s pKa, the acid is predominantly in its anionic (dissociated) form, requiring protein mediated systems to cross the plasma membrane (24,25).

The Acetate Uptake Transporter (AceTr) family (TCDB 2.A.96, Pfam: PF01184) is evolutionarily conserved across bacteria, archaea and fungi (26). A majority of fungal genomes (97 %) include members of this family, suggesting crucial roles in fungal growth and development (27,28). Key transporters of the AceTr family include Gpr1 in *Yarrowia lipolytica* (29), Ady2/Ato1 in *Saccharomyces cerevisiae*, AcpA in *Aspergillus nidulans* (30) and Satp in bacteria (31). Members of the AceTr family carry six transmembrane segments (TMS). The conserved signature motif NPAPLGL(M/F) at the beginning of the first transmembrane segment (TMS) plays a critical role in substrate uptake (26,32). The only elucidated crystal structures among AceTr family members are those of SatP from *Escherichia coli* (EcSatP) and *Citrobacter koseri* (SkSatP) (33,34). Structural analysis of SatP has confirmed six transmembrane helices, four substrate-binding sites, and a constriction site located in the center of the protein pore. This constriction site is formed by three hydrophobic residues, phenylalanine (F), tyrosine (Y), and leucine (L), known as the FLY motif (structural element) which is essential in substrate selectivity and specificity (32,33).

In *Esherichia coli*, EcSatp facilitates the transport of monocarboxylates (acetate and lactate) and dicarboxylate (succinate) (31–33). In *S. cerevisiae*, Ato1, is identified as an acetate permease with additional capability of transporting monocarboxylates, including propionate, formate and lactate (26,30). and has two paralogs: Ato2 (or Fun34) and Ato3. The expression of these three transporters is induced significantly under carbon-limiting conditions and following entry to stationary phase after growth on glucose (35,36). Palková et al., 2002 reported that the ScAto transporters (Ato1, Ato2, and Ato3) drive increases in the external pH by expelling ammonium and importing protons. The *ScATO1* gene is also required for protein-mediated acetate import, when *S. cerevisiae* cells are grown in acetate at pH 6.0 (30). Homologs of *ScATO1* have been found in *Candida* species. Indeed, twelve of the most medically relevant Candida spp. contain at least two homologous genes, and the majority have five or more. The expanded *ATO* gene families in *Candida* pathogens reinforces the idea that they play important roles in fungal adaptation and probably contribute to virulence (8,26). *C. albicans*, has a large AceTr family that includes ten *ATO* (Acetate Transporter Ortholog) genes (ATO1, ATO2, ATO3, ATO4, ATO5, ATO6, ATO7, ATO8, ATO9, and ATO10)(8). The functions of the respective proteins remain largely unknown (37). *CaATO9* and *CaATO10* are probably pseudogenes, originated from a single gene that was disrupted by insertion of transposable element (8). Previous studies have shown that the neutralization of the acidic phagolysosome induces the differentiation of *C. albicans* yeast cells to form filamentous hyphae, thereby facilitating their escape from macrophages via immune cell rupture (38). Notably, mutations in certain *C. albicans ATO* genes disrupt this process, impairing phagolysosomal neutralization, hyphal differentiation, and macrophage killing (37). In a murine model of gut colonization, deletion of the entire *ATO* gene family significantly reduces *C. albicans* colonization, particularly after antibiotic elicited bacterial disruption, indicating that *ATOs* are important for long-term persistence in the gut (Alves et al., 2025, unpublished data, manuscript under review).

In this study, we focused on the phylogenetic analysis of AceTr family members, the *in silico* structural characterization of Ato proteins divergent *Candida* species (*C. albicans, C. glabrata*, and *C. auris*), and the cellular expression of CaAto proteins. Our analysis reveals the conservation of significant AceTr specific motifs across the mentioned species alongside the presence of distinct pore radius in the respective Ato proteins, suggestive of distinct transport activities and specificities. Additionally, we show that *ATO1, ATO2*, *ATO3* and *ATO6* are differentially expressed in *C. albicans*, with *ATO1* playing a key role in regulating the expression of other Ato family members for the assimilation of short-chain carboxylic acids like acetic and lactic acids. Finally, we describe a previously unknown ATO-related family in vertebrates that exhibits ATO-like transport domains unexpectedly fused to the Sua5/YciO/YrdC domain involved in the formation of threonyl-carbamoyl-adenosine in tRNA modification.

## 2. Methods

### 2.1 *In silico* analysis

#### 2.1.1 Protein sequence alignments and phylogenetic construction

Entries containing the PFAM annotation Gpr1_Fun34_YaaH (PF01184) were collected from UniprotKB using the Uniprot API. The sequences were aligned using MAFFT-LINSI (39) and positions with >90% gaps were removed from the alignment with TrimAl (40). Trees were constructed using IQtree v2.0 (41) with model selection on auto (selected model: Q.pfam+G4) and 1000 iterations of ultrafast bootstrap. Manual inspection was performed for the fungal trees and almost identical branches (representing the same sequence from different strains of the same species or different proteins isoforms) were removed. Trees were visualized using ETE4 (https://github.com/etetoolkit/ete4) (42) and Adobe Photoshop.

The identification of the entire conserved amino acid residues of the AceTr family was performed with the Clustal Omega software (43) (www.clustal.org). The protein sequences of the following AceTr members were included in the multiple sequence alignment: *S. cerevisiae* Ato1, Ato2, Ato3; *C. albicans* Ato1, Ato2, Ato3, Ato4, Ato5, Ato6, Ato7, Ato8; *C. glabrata*: Ato1, Ato2, Ato3 and *C. auris*: Ato1, Ato2, Ato3 (Table 1).

**Table 1.** Accession number of Ato proteins of *C. albicans*, *C. glabrata*, *C. auris* and *S. cerevisiae* in NCBI and AlphaFold Databases. The “Database nomenclature” refers to the nomenclature used in NCBI. The “Alias” denotes the nomenclature recently proposed by our group (8). The Alias was used for the analysis performed in this study.

| Species | Database nomenclature | Alias (8) | Accession no. in NCBI | Accession no. in AlphaFold Database |
| --- | --- | --- | --- | --- |
| *C. albicans* | FRP3 | ATO1 | XP_710295.1 | AF-A0A1D8PHP8-F1 |
|  | ATO1 | ATO2 | XP_710650.1 | AF-A0A1D8PJ22-F1 |
|  | ATO2 | ATO3 | XP_718515.2 | AF-A0A1D8PJ42-F1 |
|  | ATO6 | ATO4 | XP_714701.1 | AF-A0A1D8PKA0-F1 |
|  | ATO5 | ATO5 | XP_714703.1 | AF-A0A1D8PKD7-F1 |
|  | FRP6 | ATO6 | XP_716748.1 | AF-Q5A4K0-F1 |
|  | FRP5 | ATO7 | XP_716747.2 | AF-A0A1D8PPM3-F1 |
|  | ATO7 | ATO8 | XP_019330752.1 | AF-A0A1D8PGM7-F1 |
|  | ATO9 | ATO10 | XP_717951.1 | AF-Q5A867-F1-v4 |
|  | ATO10 | ATO9 | XP_717953.1 | AF-Q5A865-F1-v4 |
| *C. glabrata* | ATO1 | - | XP_449497.1 | AF-Q6FJU7-F1 |
|  | ATO2 | - | XP_449115.1 | AF-Q6FKX9-F1 |
|  | ATO3 | - | XP_444907.1 | AF-Q6FYA8-F1 |
| *C. auris* | ATO1 | - | XP_028889701.1 | AF-A0A2H0ZYE7-F1 |
|  | ATO2 | - | XP_028890140.1 | AF-A0A2H0ZRA6-F1 |
|  | ATO3 | - | XP_028890274.1 | AF-A0A2H0ZM18-F1 |
| *S. cerevisiae* | ATO1 | - | NP_009936.1 | AF-A0A815X247-F1 |
|  | ATO2 | - | NP_014399.3 | AF-C7GLQ2-F1 |
|  | ATO3 | - | NP_010672.1 | AF-A0A815XMT5-F1 |

#### 2.1.2 Three-dimensional structure and pore radius predictions

The 3D structures of Ato proteins from *S. cerevisiae*, *C. albicans*, *C. glabrata* and *C. auris* were obtained from the AlphaFold Protein Structure Database (Table 1)(44,45). These structures were used to predict the radius along the pore using the HOLE program (2.2.005 Linux) (46). The HOLE program provided the Van der Waals *file simple radii*, which was used to predict the pore of each Ato protein. These predictions were visualized along with 3D protein structures by the Visual Molecular Dynamics program (VMD, 1.9.3)(47). The pore radius of the Ato proteins was presented on a graphic with the X coordinates representing the length of the Ato proteins (48).

#### 2.1.3 Molecular docking

Molecular docking simulations with acetate and lactate substrates were performed using Auto Dock Vina from PyRx virtual screening tool (49), as previously described (50). The ligand structure of acetate and lactate were obtained from PubChem (https://pubchem.ncbi.nlm.nih.gov/). The simulated interactions of ligands and proteins were visualized and analyzed by Maestro v11.2 and Chimera, a graphical user interface for AutoDock Vina (50). The dissociated forms of each carboxylic acid were used in the docking studies.

#### 2.1.4 Domain annotation

Domains were predicted using Hmmscan (HMMER 3.4, Aug 2023; http://hmmer.org/) for the HMM-profiles of interest (PF01184: Gpr1_Fun34_YaaH and PF01300: Sua5_yciO_yrdC). Hits with a bit-score > 25 for PF01300 and > 0 for PF01184 (increasing sensitivity to predict some of the more divergent metazoan domains) were considered. Domains were visualized next to the trees using ETE4.

#### 2.1.5 Structural superimposition and visualization

The crystal structure for SatP (33) (5ZUG) and an Alphafold 2.0 predicted structure for A0A3B3D9A3 (44) were used. Structures were superimposed using the TM-align webserver (https://zhanggroup.org/TM-align/ ) for TM-score and RMSD inference. For visualization PyMOL (Version 3.0 Schrödinger, LLC) was used. Structures were superimposed using the PyMOL super command and figures were finalized using Adobe Photoshop.

#### 2.1.6. Phosphorylation sites prediction

Phosphorylation site prediction was performed using Netphos-3.1b (https://services.healthtech.dtu.dk/services/NetPhos-3.1/) (51,52). This tool uses multiple neural networks to make predictions.

### 2.2 Strains and growth conditions

All *C. albicans* strains generated in this study, listed in Table 2, were derived from the wild-type (WT) clinical isolate *C. albicans* SC5314. Strains generated using the CRISPR-Cas9 system were selected on YPD medium supplemented with 200 µg/mL nourseothricin. Following the removal of the CRISPR-Cas9 cassette, the strains were cultured on yeast extract-peptone-dextrose (YPD) medium, comprising yeast extract (1% w/v), peptone (1% w/v), glucose (2% w/v), and agar (2% w/v). For microscopy analysis, cells were grown in synthetic minimal medium (SM: 0.67% w/v YNB with ammonium sulphate) containing 0.2% glucose and then transferred to synthetic minimal medium supplemented with specific carboxylic acids: acetate (0.1% v/v, pH 6.0) or lactate (0.1% v/v, pH 5.0) at 30 °C. The pH of the media containing carboxylic acids was adjusted with a 5 M NaOH solution.

**Table 2.**
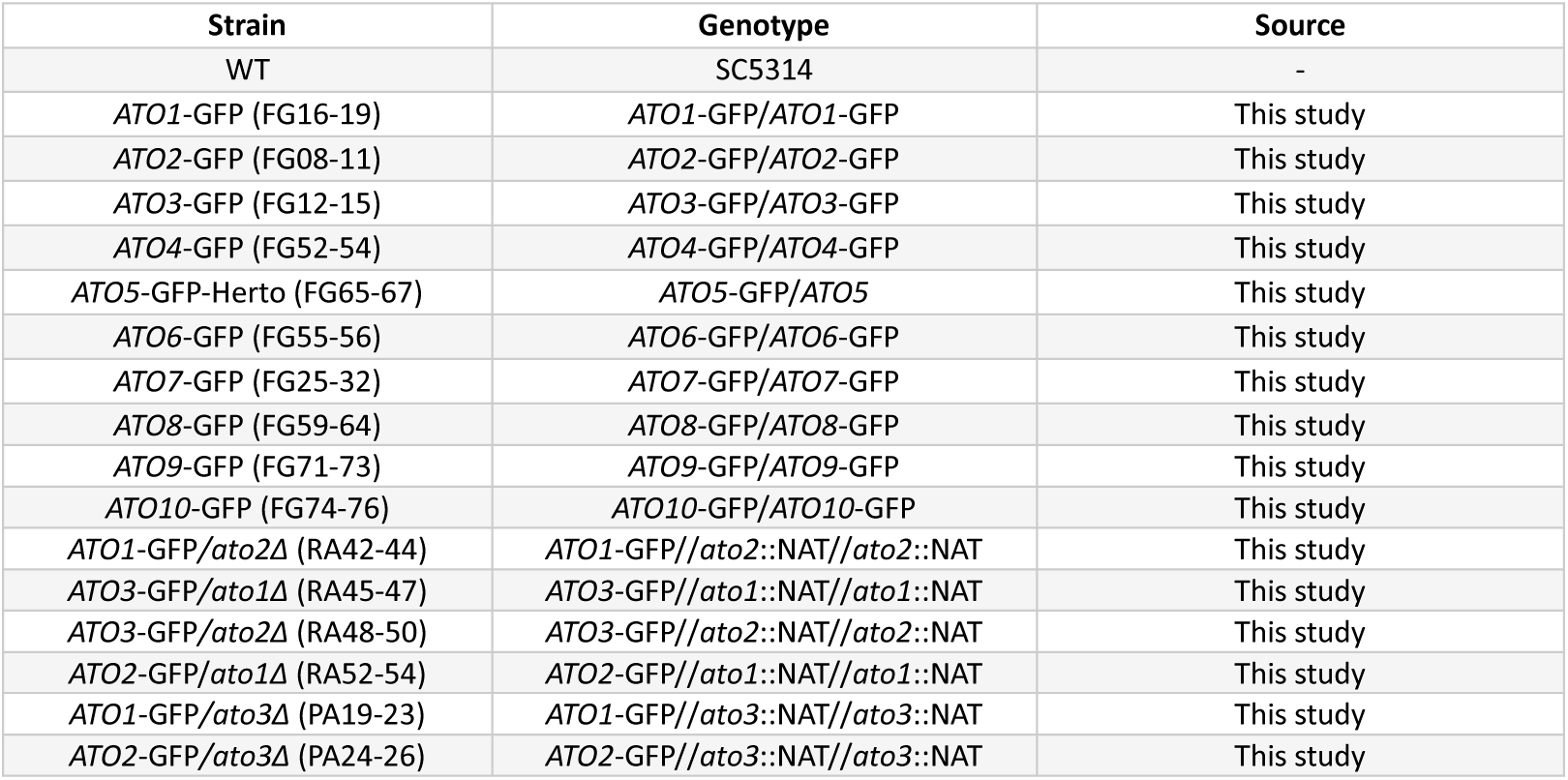
List of strains used in this study

| Strain | Genotype | Source |
| --- | --- | --- |
| WT | SC5314 | - |
| *ATO1*-GFP (FG16-19) | *ATO1*-GFP/*ATO1*-GFP | This study |
| *ATO2*-GFP (FG08-11) | *ATO2*-GFP/*ATO2*-GFP | This study |
| *ATO3*-GFP (FG12-15) | *ATO3*-GFP/*ATO3*-GFP | This study |
| *ATO4*-GFP (FG52-54) | *ATO4*-GFP/*ATO4*-GFP | This study |
| *ATO5*-GFP-Herto (FG65-67) | *ATO5*-GFP/*ATO5* | This study |
| *ATO6*-GFP (FG55-56) | *ATO6*-GFP/*ATO6*-GFP | This study |
| *ATO7*-GFP (FG25-32) | *ATO7*-GFP/*ATO7*-GFP | This study |
| *ATO8*-GFP (FG59-64) | *ATO8*-GFP/*ATO8*-GFP | This study |
| *ATO9*-GFP (FG71-73) | *ATO9*-GFP/*ATO9*-GFP | This study |
| *ATO10*-GFP (FG74-76) | *ATO10*-GFP/*ATO10*-GFP | This study |
| *ATO1*-GFP/*ato2Δ* (RA42-44) | *ATO1*-GFP//*ato2::NAT*//*ato2::NAT* | This study |
| *ATO3*-GFP/*ato1Δ* (RA45-47) | *ATO3*-GFP//*ato1::NAT*//*ato1::NAT* | This study |
| *ATO3*-GFP/*ato2Δ* (RA48-50) | *ATO3*-GFP//*ato2::NAT*//*ato2::NAT* | This study |
| *ATO2*-GFP/*ato1Δ* (RA52-54) | *ATO2*-GFP//*ato1::NAT*//*ato1::NAT* | This study |
| *ATO1*-GFP/*ato3Δ* (PA19-23) | *ATO1*-GFP//*ato3::NAT*//*ato3::NAT* | This study |
| *ATO2*-GFP/*ato3Δ* (PA24-26) | *ATO2*-GFP//*ato3::NAT*//*ato3::NAT* | This study |

### 2.3 Genetic manipulation of *C. albicans* using the CRISPR-Cas9 system for generation of GFP fusions and gene deletions

GFP fusions at the 3’ terminal of *ATO* genes were performed using the CRISPR-Cas9 system following an adapted version of the protocol from Hernday lab (53). In summary, each gene’s customized gRNA expression cassette (Fragment C) was generated by assembling fragments A and B using cloning-free stitching PCR (with primers AH01237 and AH01236). Fragment A was amplified from pADH110 (with AHO1096 and AH1098 primers), while fragment B was amplified from pADH147 (with AHO1097 and Hernday-GOI primers). The Cas9 cassette was obtained by digestion of the pADH99 plasmid with the *MssI* restriction enzyme. In a single transformation, the Cas9 and gRNA cassettes were co-transformed along with a specific donor’s DNA-GFP (linear DNA fragment), which was amplified from the template CIp10 - γmGFP (with primers dDNA-GOI-GFP-FW and dDNA-GOI-GFP-RV) (54). *C. albicans* positive transformants were selected on YPD medium supplemented with nourseothricin and verified by colony PCR (with desired gene-FW-check and desired gene-RV-check or Inside-GFP-Check-RV primers). After confirming GFP integration, the NAT marker, Cas9, and gRNA expression cassettes were removed by inducing the maltose- inducible FLP recombinase system in YP medium with 3% maltose, followed by overnight incubation at 30°C. To verify the effective removal of CRISPR components, cells were streaked onto YPD agar to isolate single colonies. They were then tested on both YPD and YPD supplemented with nourseothricin. Next, all generated strains were further tested by colony PCR and confirmed through sequencing. *C. albicans ATO- GFP* strains, carrying a single *ato* knockout, were generated using a modified transient CRISPR-Cas9 approach (55). All sgRNA expression cassettes were constructed through a three-step PCR process using the pV1093 plasmid as a template. The steps included (i) synthesizing the SNR52 promoter (with oligonucleotides: SNR52/F and SNR52/R-desired gene), (ii) generating the gRNA scaffold (with primers: sgRNA/F-desired gene, and sgRNA/R) and (iii) fusing the SNR52 promoter with gRNA scaffold cassettes (with oligonucleotides: SNR52/N and sgRNA/N). The Cas9 expression cassette was amplified from the pV1093 plasmid via PCR using the CaCas9/F and CaCas9/R primers. Similarly, repair templates specific to the desired gene were generated through PCR using the primers NAT-desired gene-repair/F and NAT- desired gene-repair/R. *C. albicans* cells were simultaneously transformed with the three constructed cassettes using the classical lithium acetate method (56). Positive *C. albicans* transformants were selected in YPD supplemented with nourseothricin and, subsequently, validated through colony PCR (using the specific oligonucleotides gene-FW-check and desired gene-RV-check). All primers used in this study are listed in Table S1.

### 2.4 Epifluorescence microscopy

*C. albicans* cells expressing Atos-GFP were grown in a synthetic minimal medium supplemented with 0.2% glucose until the exponential phase at 30 °C, 200 rpm. Cells were then washed and transferred to synthetic minimal media containing acetate (0.1% v/v, pH 6.0), lactate (0.1% v/v, pH 5.0), or without carbon source at 30 °C, 200 rpm. Samples were collected at 0 hours, 5 hours, and 24 hours after derepression for analysis by epifluorescence microscopy and for further protein extraction (see section 2.5). For microscopy analysis, 1 mL of each culture was harvested, cells were concentrated by centrifugation and manually immobilized on slides. Cells were visualized using a Leica DM5000B microscope with appropriate filters. Images were captured with a Leica DFC 350FX R2 digital camera using LAS AF V1.4.1 software (Leica). The images presented are representative of three independent experiments.

### 2.5 Western Blot and quantification analysis

Yeast cells were grown as explained above in section 2.4. At the time-point 5 hours, cells were harvested at a final OD_640_ of 1.5 per mL, and total protein extracts were prepared using the NaOH/TCA method (57). Briefly, cells were harvested by centrifugation (max. rpm, 1 min, 4 °C) and the pellet was resuspended in 500 µL of cold water. Cell lysates were prepared by adding 50 µL of NaOH (1.85 M) following 10 min of incubation on ice. Then, 50 µL of trichloroacetic acid (TCA, 50%) was added and the samples were incubated for an additional 10 min on ice. Protein precipitates were collected by centrifugation (max. rpm, 15 min, 4 °C). The pellets were dissolved in 50 µL of loading buffer (33.3 mM Tris-hydrochloride pH 6.8; 1.33 mM EDTA; 1.33% sodium dodecyl sulphate (SDS); 6.66% glycerol; 0.02% bromophenol blue and 2% Beta-mercaptoethanol) and heated at 37 °C for 10 min. Each sample (10 µL) was loaded onto a 10% acrylamide gel. Following electrophoresis, the separated proteins were transferred onto nitrocellulose membranes (GE Healthcare Life Sciences. For transference and loading controls, the membranes were stained with a Ponceau S solution (0.1% Ponceau (w/v); 0.5% glacial acetic acid (v/v)). Following washing and blocking steps, the membranes were probed with the primary antibody anti-GFP (monoclonal, mouse IgG1κ, clones 7.1 and 13.1, Roche, 11814460001) used at 1:3000 dilution. The antimouse-IgG (whole molecule)-peroxidase produced in rabbit (A9044, Sigma) was used as a secondary antibody at 1:100000 dilution. The signal was detected by enhanced chemiluminescence using the WesternBright ECL HRP substrate (advansta). All images were captured using a Syngene G:Box Chemi XX9 image documentation system and GeneSys software. The quantification levels of the GFP signal were performed using the ImageJ software (version 1.53k). The data is represented in arbitrary units (A.U.) as the mean of at least three independent experiments (n≥3) (Figure S8). The error bars represent the standard error of the mean (SEM) (Prism 8.0; GraphPad software, version 8.0.1).

## 3. Results and Discussion

### 3.1 *C. albicans* Ato Members possess conserved NPAPLGL, FLY and SY[F]GFW motifs

Multiple sequence alignment of AceTr members from *S. cerevisiae*, *C. albicans*, *C. glabrata* and *C. auris* showed that CaAto1-8, CgAto1-3, CauAto1-3 and ScAto1-3 contain common conserved regions (Figure 1). *C. albicans* Ato9 and Ato10 were excluded from this analysis as they are likely functionally inactive (8). However, manually fusing CaAto9 (2 TMS near the N-terminus) and CaAto10 (4 TMS near the C-terminus) sequences does yield a structure similar to other *C. albicans* Atos (Figure S1) which may suggest that they represent fragments of one single gene. Notably, the putative functional N^88^PAPLGL^94^ signature motif (numbering refers to CaAto-1), which is localized at the beginning of the first TMS (Figure S2), is fully conserved in CaAto1-3, CgAto1-2, CauAto2, and ScAto1-2 (Figure 1). In CaAto6-8 and CauAto3, the first Pro residue in this motif (Pro89) is substituted by Ala or Ser. In addition, in CaAto4-7 and CauAto3, the second Pro residue (Pro91) is substituted by Ala. The highly conserved residue Leu92 is substituted by isoleucine in CaAto4 and CaAto5 and valine in CaAto7. Therefore, CaAto7 and CaAto8 exhibit the highest divergence in the NPAPLGL motif of these ATO families, differing in four and three of the sixteen residues in this motif, respectively. These differences are likely to change the substrate affinities, specificities and/or activities of these transporters. This prediction is based on previous mutational analyses of ScAto1, which established that amino acid substitutions in this motif impaired protein function *per se*, without affecting its proper localization to the plasma membrane (50).

**Figure 1.**
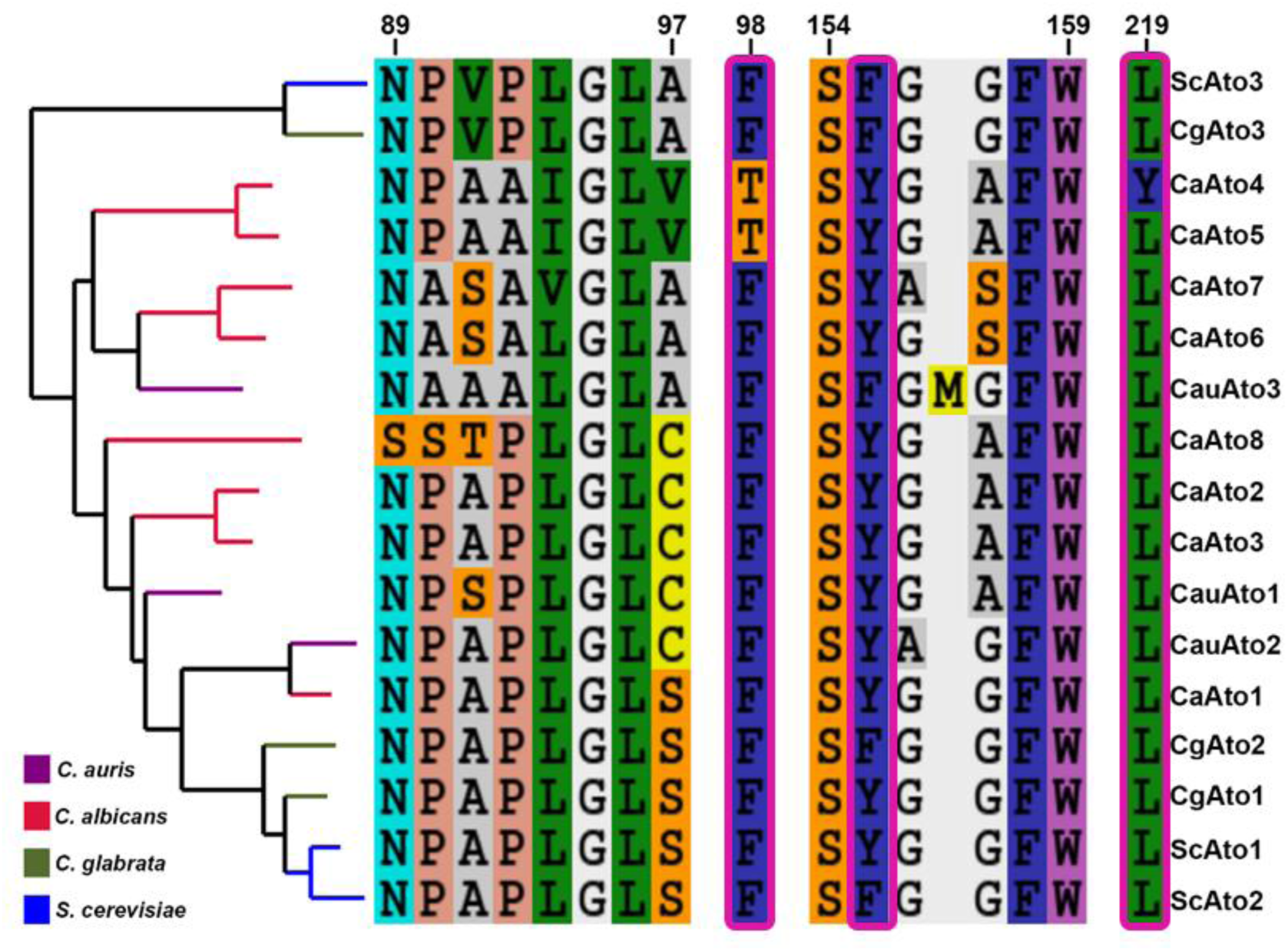
Phylogeny and important positions of Ato homologous sequences from *C. albicans* (Ato1-8), *C. glabrata* (Ato1-3), C. auris (Ato1-3) and *S. cerevisiae* (Ato1-3). The signature motif, N-P-A-P-L-G-L of the AceTr family (58), the narrowest hydrophobic constriction region (FLY) (pink rectangles) (33) and the signature motif S-Y[F]-G-F-W are aligned next to the tree. Full alignment is available in the supplementary data (Figure. S2).

In ScAto1, the narrowest constriction site is formed by the FLY structural element. Previous studies have demonstrated that mutations of FLY residues of ScAto1 lead to altered protein activity and specificity, highlighted by the observation that Leu219Ala allows cells to grow on succinic acid (32). All analyzed *Candida* homologs possess a well-conserved phenylalanine in this FLY motif (Phe98, Figure 1), with the exception of CaAto4 and CaAto5, where it has been substituted by threonine. The substitution of a bulky hydrophobic residue by a polar amino acid might influence substrate transport through the inner pore of the transporter, as previously described for ScAto1 (32). The Tyr residue of the FLY motif is also highly conserved, being replaced by phenylalanine, a residue with similar biochemical character, in ScAto2, ScAto3, CgAto2, CgAto3, and CauAto3 (Figure 1). Finally, the Leu residue of the FLY motif is nearly absolutely conserved with the only exception of CaAto4, where it is a Tyr. Also, the Tyr residue of the FLY structural motif is also part of the SY[F]GFW linear signature motif of the AceTr family, located in TMS3 (Figure S2). This motif is also highly conserved among the Ato proteins examined. Overall, our *analysis* suggests that conserved evolutionary substitutions in the NPAPLGL, FLY and SY[F]GFW motifs might have resulted in modulations of transport activity and/or substrate specificity of Ato proteins in these *Candida* pathogens.

### 3.2 *C. albicans* Atos have distinct pore radius

The pore radii of Ato proteins in *C. albicans*, *C. glabrata* and *C. auris* were predicted to determine the putative translocation paths for their substrates (e.g. acetate or lactate) (Figure 2). The narrowest constriction in each path is located around the FLY region (Figure 2A). In CaAto4 the pore seems to be blocked in this region, due to the existing substitutions in the FLY motif. Additionally, CaAto4 includes an additional restriction site located in the upper region of the protein, that faces the extracellular medium, specifically near the residues Phe94, Phe150, and Phe218. Also, the central pore of CaAto7 appears blocked despite the conservation of FLY motif. This blockage may be caused by variations in the amino acid residues that interact with the closet transmembrane segment (TMS). On the other hand, there is a widening of the pore around the FLY motif of CaAto5 due to a substitution to TLY in this protein. In fact, the narrowest constriction site of CaAto5 is found in the upper section of the protein facing the extracellular space (Phe93, Phe149, and Phe217), similar to the second constriction site of CaAto4. An additional constriction to the FLY region has been predicted for CgAto2 and CauAto1, closer to the extracellular region (Figure S3, A, B).

**Figure 2.**
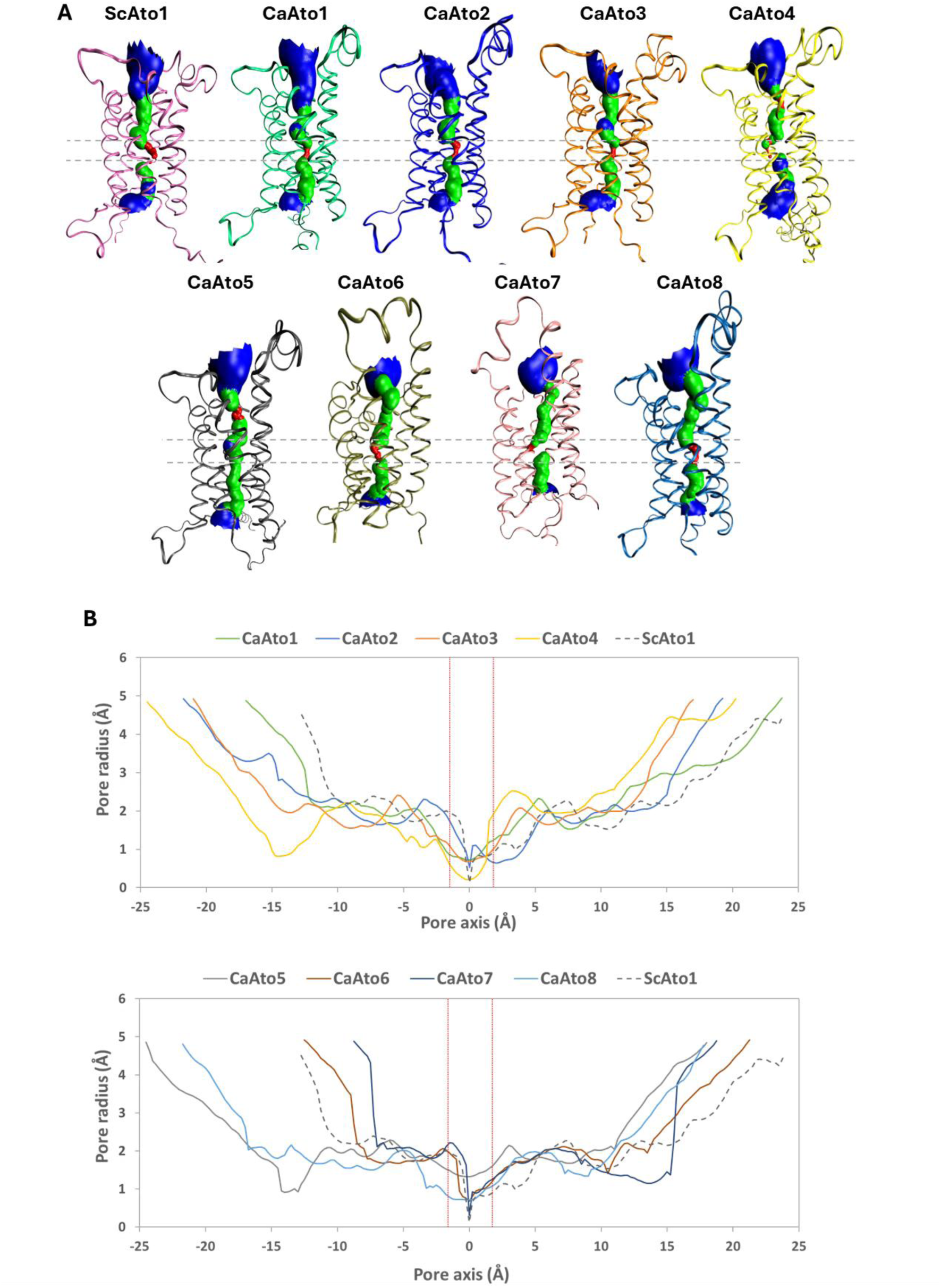
Pore 3D structure predictions and radius profiles simulations along the channel axis in the Atos of *S. cerevisiae* and ***C. albicans*. A.** Pore 3D structure prediction of ScAto1 and CaAto1-8. The 3D structures of proteins are represented in New Ribbons, with the color scheme consists of blue (representing a bigger pore size), green (representing an intermediate pore size), and red (representing a more constricted pore size) in pore prediction. The horizontal dashed grey lines correspond to the constricted site where the pore radius is tight. **B.** Simulations for the pore radius profiles along the channel axis in the ScAto1 and CaAto1-8. The region in the middle of proteins that contains the construction site is marked by vertical red dashed lines.

The variations in pore radius between CaAto4, CaAto5, CaAto7 and CaAto8 compared to ScAto1 and CaAto1(Figure 2B) suggest that these proteins might possess altered transport activity or specificity compared to ScAto1 or CaAto1. The reduction of the pore radius in CaAto4 and CaAto7 might result in the uptake of only small carboxylic acids such as formate or, or is suggestive of a signaling rather than transport role. Indeed, Rendulic and colleagues (2021) found that the substitution of Leu219 with tryptophan (L219W) in the Ato1 mutant *S. cerevisiae* strain led to a reduction of the pore radius at the construction site thereby blocking transport of carboxylic substrates. Therefore, a larger side-chain at this position can arrest transport. Furthermore, the pore radius of CaAto5 in the FLY motif region is wider compared to ScAto1 and CaAto1, suggesting that CaAto5 may be capable of transporting larger molecules, such as succinate and citrate. Indeed, Phe98Ala and Leu219Ala mutations in ScAto1 reduce the constriction at the central FLY region and enable the transport of larger molecules, such as succinate (32). In addition, Tyr155Phe and Leu219Val substitutions in the FLY motif of *Candida jadinii* Ato5, a broad range carboxylate transporter that transports mono- di and tricarboxylates, including citrate (58). These findings indicate that constrictive FLY amino acids play a crucial role in the substrate selectivity and transport activity of Ato transporters.

We also identified significant variations between CaAto5, CaAto4 and CaAto7 in their pore radii around the NPAPLGL(M/F) and SYG(X)FW motifs (Figure 1, 2). Also, the pore size prediction displays distinct dissimilarities in the pore radius of CaAto8, not only in the constriction site, but also in the conservation motif region (Figure 2B), which is less well conserved in CaAto8 (Figure 1). On the other hand, given the similarities in structure, pore size and conserved motifs, CaAto1, 2, 3, 6 may transport similar substrates to ScAto1.

The pore size analysis of the *C. glabrata* Atos shows that CgAto2 contains two distinct constriction sites (Figure S3, A, C). No significant variation in pore radius was observed between the Atos of *C. glabrata*. However, significant differences in pore radius were identified between *C. auris* CauAto1-2 and CauAto3 (Figure S3, D). The extracellular loop regions that link transmembrane segments in CauAto3 are significantly shorter than those in CauAto1-2, leading to dissimilarity in their 3D structures and pore sizes. In addition, the pore radius analysis shows two constriction sites for CauAto1 (Figure S3, B).

*S. cerevisiae* strains harbouring wild-type *ATO3* and *ATO2* genes are unable to grow on lactic acid, whereas *ATO2^T653C^* and *ATO3^T284C^* mutants can grow on this carboxylic acid (59). Therefore, enlarging the hydrophobic constrictions in these Ato transporters appears to have empowered lactate assimilation by *S. cerevisiae*. Increased binding affinity may also have contributed to an enhanced transport capacity by facilitating ligand passage through the structure. Amino acid residues outside the constriction pore may also have increased transport capacity, for example by stabilising the transporter in the plasma membrane or enhancing transitions between its closed and open states.

### 3.3 Acetate and lactate are potential substrates for *C. albicans* Ato1, Ato2, Ato3 and Ato6

The crystal structures of EcSatP and CkSatP indicate a hexameric anion channel (33,34). Each monomer comprises six transmembrane helices and four acetate binding sites aligned along the pore. Acetate binding site S1 faces the cytoplasm, sites S2 and S3 are located inside the main pore surrounding the FLY motif, and site S4 faces the periplasm (34). The FLY motif is located centrally in the pore of the EcSatP monomer, between S2 and S3 sites, leading to the formation of an hourglass-shaped structure.

We performed substrate docking with acetate and lactate for Ato proteins of *S. cerevisiae*, *C. albicans*, *C. glabrata* and *C. auris* Atos using Dock Vina from PyRx virtual screening tool. This revealed at least four predicted binding sites along the channel axis of all proteins we tested (Figure 3 and Figure S4). This was consistent with the crystal structures of the bacterial homologs EcSatP (33) and SkSatP (34). The number and positions of predicted substrate binding sites were similar for all proteins, except for CaAto4, 5, 7, 8 which contained fewer binding sites and theoretically altered binding affinities for acetate and lactate. No acetate and lactate binding sites were identified in the upper region of the CaAto5, suggesting that CaAto5 might not transport acetate or lactate. The molecular docking results for CauAto1 and CauAto3 indicate fewer and distinct binding sites for acetate and lactate, whereas CauAto2 exhibits binding sites nearly identical to other Atos (Figure S4, B).

**Figure 3.**
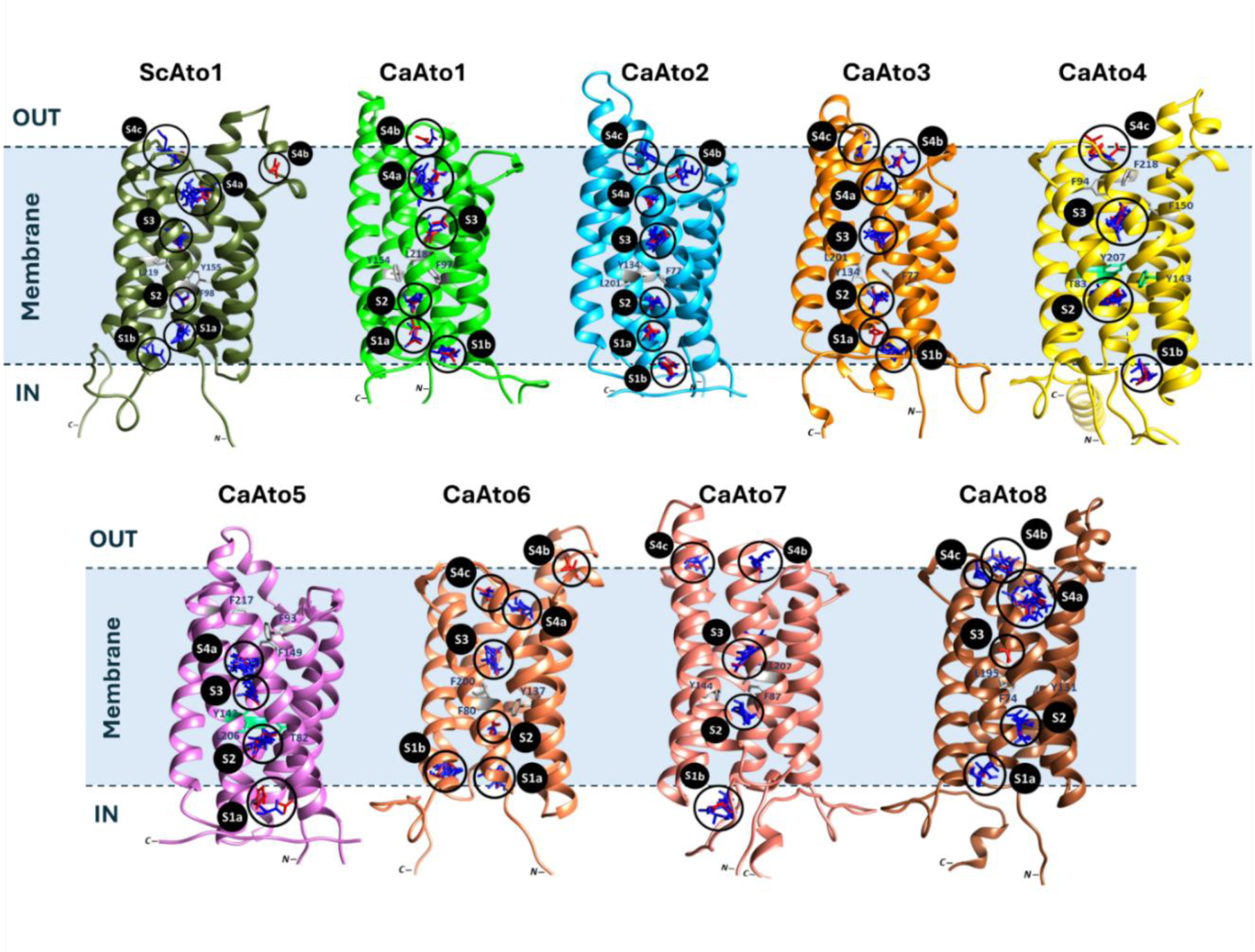
3D structure and molecular docking of ScAto1 and CaAto1-8 with acetate and lactate as substrates. Predicted binding sites for acetate and lactate were shown with S1 to S4. Site S1 is located at the cytoplasmic vestibule, Sites S2 and S3 are located inside the main pore, and Site S4 is located at the periplasmic vestibule. Localization of the N- and C-terminal of the proteins is shown. Acetate and lactate ligands are presented in red and blue respectively.

### 3.4 Specific *C. albicans* Atos are localized to the plasma membrane in the presence of acetate and lactate

To study their expression and sub-cellular localization, we generated C-terminal GFP fusions for all ten Ato proteins in *C. albicans* (Table 2). The expression of these Ato-GFP fusions was examined during growth in the absence (YNB) or presence of lactic or acetic acid. Only Ato1-GFP, Ato2-GFP, Ato3-GFP and Ato6- GFP were expressed at detectable levels (Figure 4A, B and C). Ato6-GFP was detectable in the plasma membrane (PM) after prolonged growth on lactate (24 h). Ato1-GFP, Ato2-GFP and Ato3-GFP were detected in PM after 5hof derepression in acetic or lactic acid as well as in the YNB control (Figure 4A, B and C). After 5h, Ato1-GFP and Ato2-GFP signals in PM were similar, while Ato3-GFP was expressed at lower levels under all of the conditions tested. After 24 h, Ato1-GFP and Ato2-GFP increased in the PM, as did the amount of fluorescence associated with the vacuolar lumen, suggesting a rise in the turnover of these proteins. At 24 h, Ato3-GFP was localized mainly in the vacuole.

**Figure 4.**
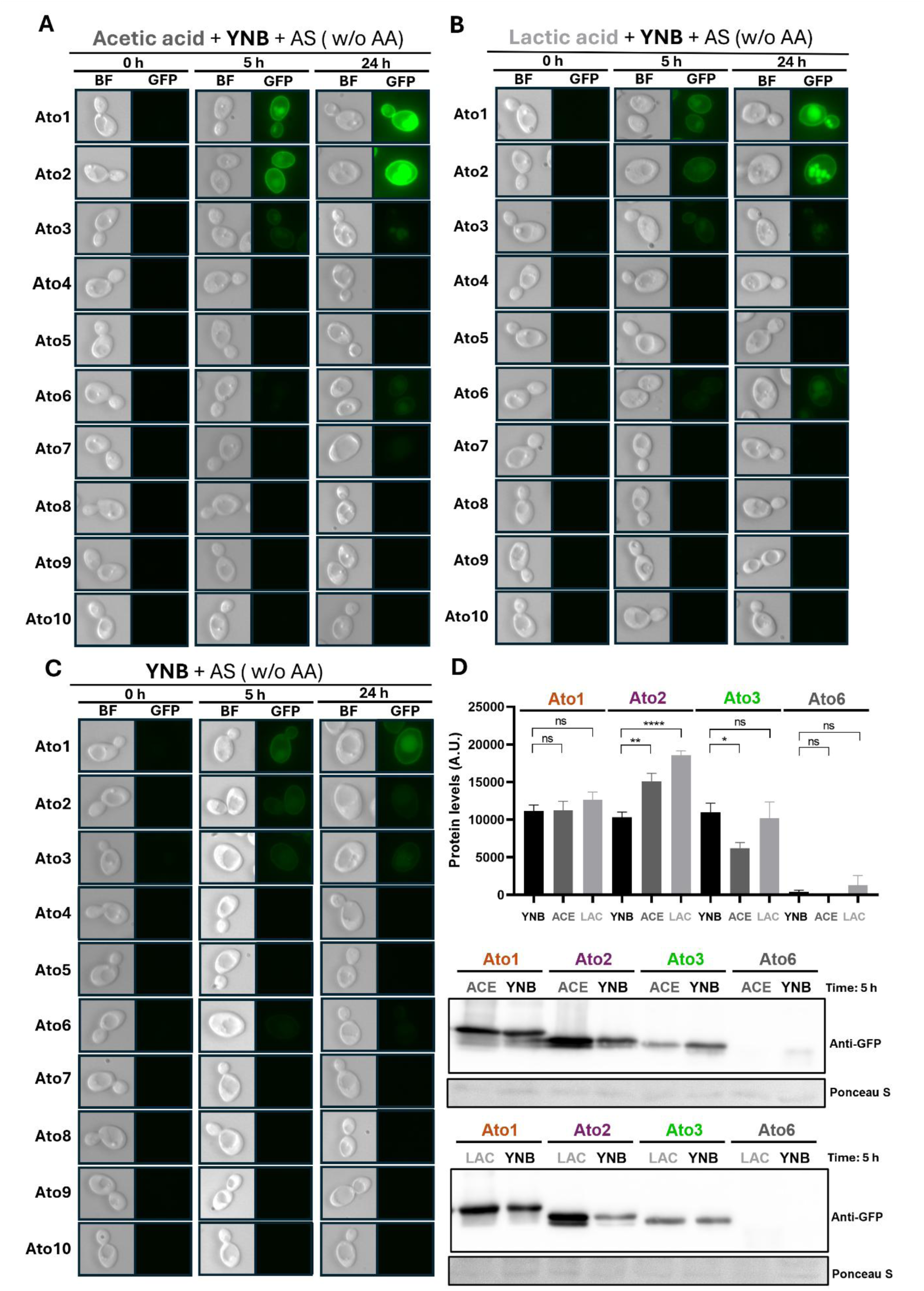
Subcellular localization of Atos-GFP proteins in live *C. albicans* cells grown on different carbon sources. *C. albicans* cells grown in 50 mL SM medium supplemented with 0.2 % (w/v) glucose to exponential phase. They were washed then with deionized water and then transferred, to fresh minimal media containing different carbon sources: (**A**) 0.1 % (v/v) acetic acid pH 6.0, (**B**) 0.1 % (v/v) lactic acid pH 5.0 and (**C**) without any carbon source (SM: 0.67% w/v YNB with ammonium sulphate). Samples were collected after 0-, 5- and 24-hours induction, to examine cells by epifluorescence microscopy. Cell cultures were grown with aeration 200 rpm, at 30 °C (**D**) At the time-point 5h, protein extracts were prepared for western immunoblotting with an anti- GFP antibody. Quantification of the protein levels in arbitrary units (A.U.) were done using ImageJ software (version 1.53k). Error bars correspond to standard error of the mean (SEM) of at least three independent experiments. ns, not significant; *P=0.0255; **P=0.0014; ****P<0.0001 (unpaired t-test (two-tailed) using Prism 8). Ponceau S staining was used as a transference and loading control. The abbreviations BF and GFP refer to "Bright-Field" and "Green Fluorescent Protein," respectively

To clarify the roles of the lactate and acetate in the expression levels of Ato1, Ato2, Ato3 and Ato6, Western-blot analyses using total protein extracts were performed (Figure 4D). Quantification of the protein levels revealed that neither acetate nor lactate significantly changed the amount of Ato1. This was rather surprising given that we have directly demonstrated that Ato1 is the main transporter responsible for radiolabeled acetate uptake (Alves et al., 2025, unpublished data, manuscript under review). On the other hand, carboxylic acids induced a significant increase in the expression of Ato2 (Figure 4D), suggesting that besides Ato1, Ato2 might play a critical role in the uptake or utilization of weak organic acids. Acetate also led to a decrease in Ato3 levels, but lactate did not change the Ato3 expression compared to the YNB control condition (Figure 4D). Ato6, showed low levels of expression at 5h with no considerable differences between the tested conditions. No signals were observed for Ato4-GFP, Ato5-GFP, Ato7-GFP, Ato8-GFP, Ato9-GFP and Ato10-GFP in response to acetate or lactate (Figure 4). This suggests that these transporters might be involved in the uptake of different substrates or be expressed in response to other stimuli. Noticeably, our *in-silico* analysis revealed no appropriate lactate or acetate binding sites in these proteins (Figure 3).

### 3.5 Deletion of *ATO1* affects the localization and expression of Ato2 and Ato3

To explore potential functional cooperative interactions between Ato proteins, we constructed *ato1, ato2* or *ato3* deletions in the Ato1- Ato2- or Ato3-GFP backgrounds in *C. albicans* (Table 2). The expression and subcellar localization of Ato-GFP proteins in these strains were examined in response to acetate or lactate (Figure 5).

**Figure 5.**
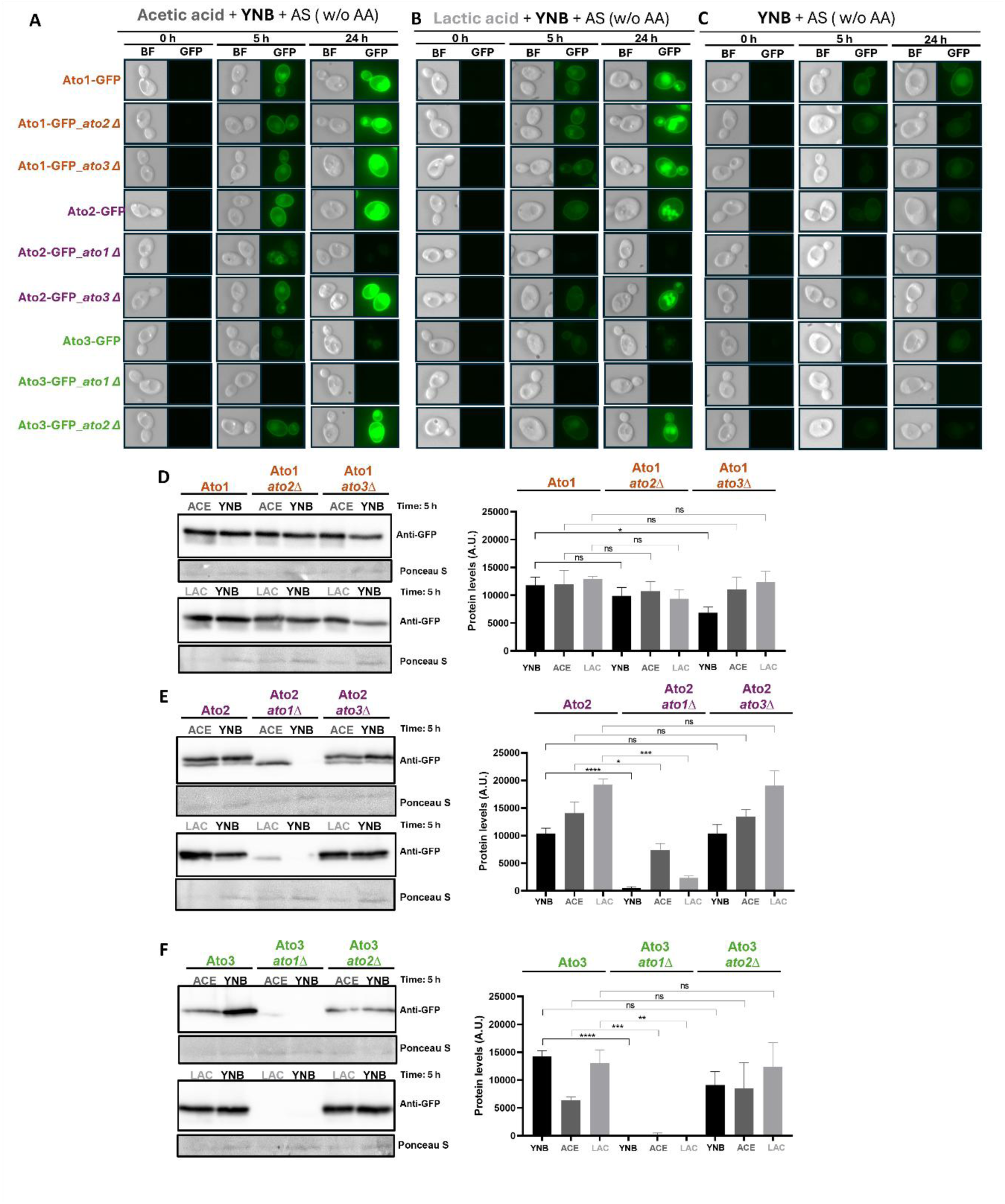
Subcellular localization of the indicted Atos-GFP in *C. albicans*. WT and *ato*-mutant cells. *C. albicans* cells grown in 50 mL SM medium supplemented with 0.2 % (w/v) glucose to exponential phase. They were washed then with deionized water and then transferred, to fresh minimal media containing different carbon sources: **(A)** 0.1 % (v/v) acetic acid pH 6.0, **(B)** 0.1 % (v/v) lactic acid pH 5.0 and **(C)** without any carbon source (SM: 0.67% w/v YNB with ammonium sulphate). Samples were collected after 0-, 5- and 24-hours induction, to examine cells by epifluorescence microscopy. Cell cultures were grown with aeration 200 rpm, at 30 °C (**D-F**) At the time-point 5h, protein extracts were prepared for western immunoblotting with an anti-GFP antibody. Quantification of the protein levels in arbitrary units (A.U.) were done using ImageJ software (version 1.53k). Error bars correspond to standard error of the mean (SEM) of at least three independent experiments. ns, not significant; *P=0.0183-0.0455; **P=0.0014-0.0054; ***P=0.0001-0.0008; ****P<0.0001 (unpaired t-test (two-tailed) using Prism 8). Ponceau S staining was used as a transference and loading control. The abbreviations BF and GFP refer to "Bright-Field" and "Green Fluorescent Protein," respectively.

The deletion of *ATO2* did not interfere with Ato1-GFP or Ato3-GFP expression under any of the conditions tested as demonstrated by GFP fluorescence (Figure 5A, B and C) and Western blotting (Figure 5D). However, deleting *ATO3* had a slight impact in expression of Ato1 only in YNB condition (Figure 5D), but did not show any influence in Ato2 protein levels in all the conditions tested (Figure 5F). Most interestingly, under all of the conditions tested, the deletion of *ATO1* had a pronounced impact on the expression and PM localization of Ato2-GFP and Ato3-GFP (Figure 5A, B and C, rows 5 and 8). The *ato1Δ* mutant exhibited Ato2-GFP localization at the cortical and nuclear endoplasmic reticulum (ER) after 5h, followed by complete abolishment of Ato2 expression after 24 hours in the presence of acetic acid. Moreover, the deletion of *ATO1* resulted in a complete loss of Ato2-GFP and Ato3-GFP expression in YNB and in response to lactate (Figure 5, B and C, lines 5 and 8). These results were consistent with the Western blot analysis, which showed either no detectable bands or only faint bands for both Ato2 and Ato3 under these conditions. Also, acetate led to a decrease in Ato3 levels, and lactate led to a slight increase in Ato6 levels, but these changes were not statistically significant. In addition, Western blotting revealed interesting effects upon Ato2-GFP expression in the presence of acetate and lactate (Figure 5E). In the wild-type background, Ato2-GFP appeared as two bands, possibly representing non-phosphorylated and phosphorylated versions of Ato2. However, in the *ato1* mutant, only the lower band was observed in response to acetate or lactate (Figure 5E). This suggests that Ato1 might regulate Ato2 by controlling (directly or indirectly) its post-translation modification. Indeed, Ato2 contains 20 potential phosphorylation sites including 12 serine and 8 threonine residues (Figure S9). Four serine residues at positions 6, 9, 13, and 28 showed approximately maximum scores (1.000), suggesting a very high probability of phosphorylation. Importantly, these residues are localized within the N-terminal region of the protein. Several phosphorylation sites have been reported in the N-terminal region of Ato proteins in *S. cerevisiae* (60). These findings reinforce that CaAto2 potentially undergoes phosphorylation. Noticeably also, after 5h in acetate, the presence of the lower Ato2-GFP band correlated with fluorescence in the endoplasmic reticulum (Figure 5A, line 5), suggesting that unmodified Ato2-GFP fails to be targeted to the PM and that this modification probably occurs at ER. In the absence of Ato1, neither form of Ato2-GFP was observed without acetate or lactate (Figure 5E, YNB control), reflecting the lack of fluorescence under these conditions at 24 h (Figure 5, B and C, line 5). The same was true for Ato3-GFP in *ato1* cells (Figure 5, B and C, line 8; Figure 5F). Overall, Ato1 appears to be required for stable expression and proper localization of Ato2 and Ato3 in *C. albicans*. This would be compatible with physical interactions between Ato1, Ato2 and Ato3 and possibly their formation of heterohexameric complexes in *C. albicans.* Some AceTr family members are thought to form homohexamers: ecSatP and ckSatP (33,34) and possibly ScAto1, ScAto2 and ScAto3 (61,62). However, there is evidence suggesting that ScAto1 and ScAto2 interact physically at the plasma membrane (57).

### 3.6 Vertebrate ATOs are characterized by transport domain duplications and a gene fusion event

Taxonomic distribution studies showed that AceTr/Ato proteins are present in both prokaryotic and eukaryotic members (26,58), but not in metazoan. These studies employed BLASTp searches, using known Ato proteins as baits (EcSatP from *E. coli,* AceP from *M. acetivorans,* Ady2 from *S. cerevisiae,* or Atos from *Cyberlindnera jadinii*). To further investigate, we used a sensitive hmmsearch and the PFAM HMM-profile (PF01184) of the AceTr transporter family to collect fungal and metazoan sequences. This revealed that AceTr/Ato family members are widespread within *Bilateria* and particularly vertebrates (Figure 6 and Figure S7). Vertebrate ATOs form two clades with representatives from every major lineage, except mammals (Figure 6). A small number of protostome sequences can be identified within the vertebrate clade, suggesting possible horizontal gene transfer. Notably, vertebrate ATOs and their close invertebrate homologs are *circa* 900 amino acids in length, contrasting with the 200-300 amino acid length of their fungal and prokaryotic counterparts (Figure 1) (33,34). Surprisingly, almost all of the vertebrate sequences that we identified have a C-terminal fusion to a Sua5/YciO/YrdC domain in addition to the Gpr1/Fun34/YaaH domains that characterize the AceTr/Ato transporter family (Figure 6 and Figure S6). This domain is not present in the bacterial SatP (Figure 6) or in the protostome and fungal clades (Figure S7). Sua5/YciO/YrdC proteins are annotated as threonyl-carbamoyl-AMP synthetases, which attach a threonyl-carbamoyl group to A37 (t6A37) of a tRNA (63). This tRNA modification is common for tRNAs that have A/U-rich anticodons, helping to stabilize their codon-anticodon interactions (64). Loss of threonyl-carbamoyl-AMP synthetase (Sua5) leads to increased rates of translational frameshifting and stop codon read-through in yeast, underlining the importance of the t6A37 modification for translational fidelity (65).

**Figure 6.**
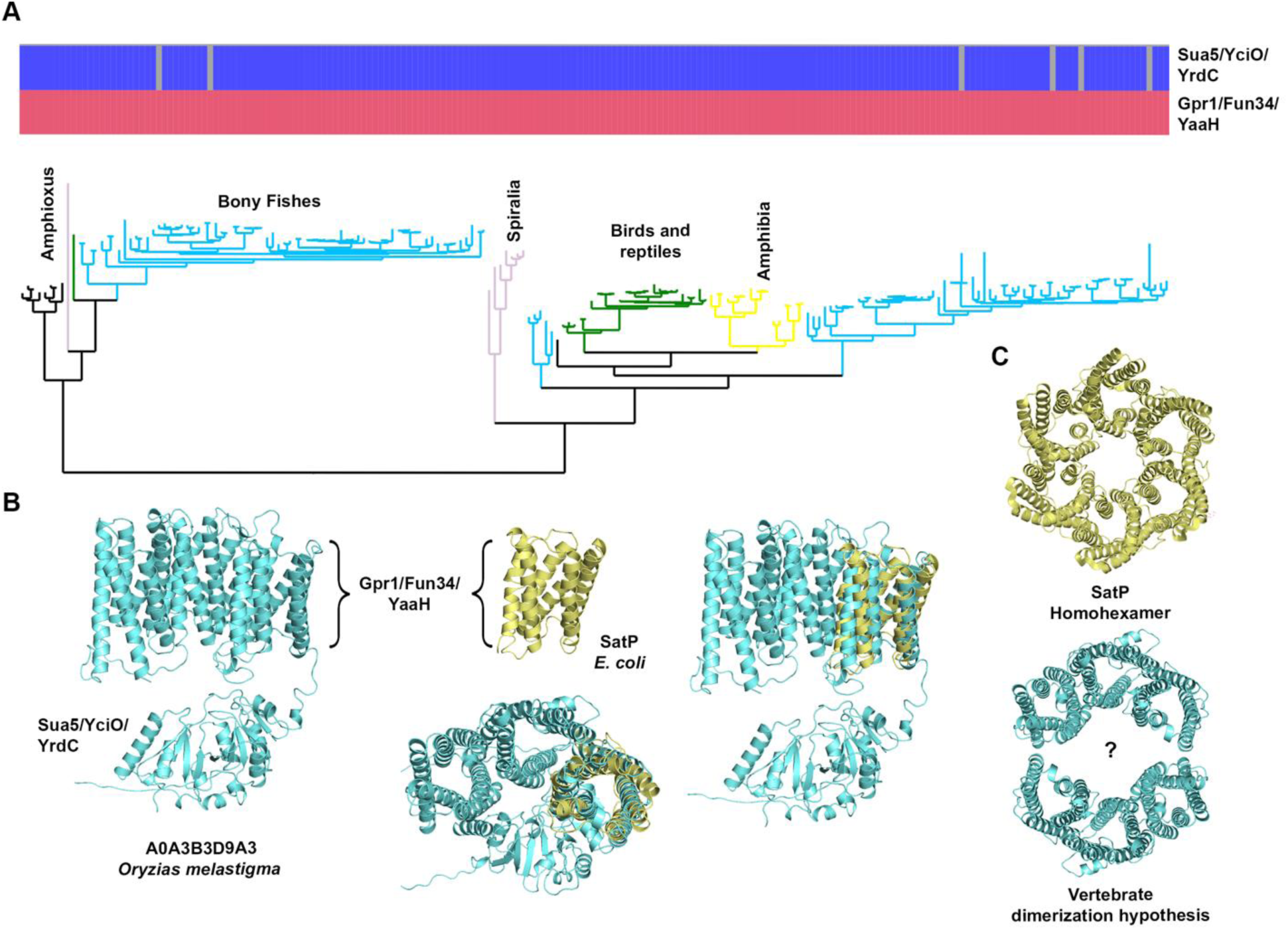
Vertebrate ATO phylogeny and structure. **A.** Phylogeny and domain composition of vertebrate ATO proteins. Domain presence is denoted as blue or red and domain absence with grey. Tree branches are colored based on taxonomy. Full metazoan phylogeny can be seen in figure S6. **B.** Alphafold 2.0 structure prediction of a vertebrate ATO protein (cyan) and crystal structure of SatP (yellow). The triplication of the Gpr1/Fun34/YaaH domain is apparent when structures are superimposed. **C.** Dimerization hypothesis of the vertebrate ATO homologs based on the experimentally determined SatP hexamer_._

Comparing the experimentally resolved EcSatP structure from *E. coli* (33) with the AlphaFold 2.0 (44) predictions of some vertebrate ATO proteins revealed that they consist of three Gpr1/Fun34/YaaH domains (six helices each) fused in tandem (Figure 6B). HMM-scan for the Gpr1/Fun34/YaaH HMM-profile in vertebrate sequences returns three domain hits for some of them (Figure S6). Structural investigation of proteins that produce less than three hits revealed that they possess three copies of the six-helix Gpr1/Fun34/YaaH fold (Figure S6).

TM-align superposition of the vertebrate A0A3B3D9A3 from *Oryzias melastigma* (chordate fish) with SatP revealed strong structural similarity with an RMSD of 2.53 Å and a TM-score of 0.82. Superimposed structures of the two proteins are presented in Figure 6. Interestingly, SatP crystals have revealed a homohexameric structure (33) (Figure 6C). Observing the duplicated Gpr1/Fun34/YaaH domains of vertebrate Atos, we hypothesized that two interacting protomers could be arranged in a six-domain ring similar to the one formed by SatP. Hence, cross-kingdom conservation of the six-domain ring structure of Ato proteins suggests that its functional relevance persists in vertebrate homologs.

## 4. Conclusion

Our study investigated the expansion, structural divergence, and functional specialization of the ATO protein family in *Candida species*, with a focus on *C. albicans*. We found that key motifs such as NPAPLGL and FLY are conserved among Ato homologs. However, significant variations exist in the pore constrictions, which are potentially responsible for their substrate specificity. Our findings revealed that Ato2 expression is markedly induced under acetate and lactate growth conditions, with prominent localization at the plasma membrane. Additionally, the expression of Ato2 and Ato3 at the plasma membrane was dependent upon Ato1, which is consistent with the idea that functional Ato proteins assemble as heterooligomeric complexes in *C. albicans*. This is further supported by the observation that Ato1 and Ato2 interact physically in *S. cerevisiae* (61,62). In addition, the bacterial homolog SatP has been reported to function as a homohexamer (33). This suggests that oligomerization may be a conserved feature required for the proper activity of these transporter proteins, whether through homomeric or heteromeric assembly. Our *in-silico* and expression analysis suggested that Ato4, Ato5, Ato7 and Ato8 are not involved in the utilization of lactate or acetate and therefore may have different substrate specificities. Our findings regarding the fusion of Ato members with Sua5 enzyme in vertebrates remain elusive. We hypothesize that the cytoplasmic region of vertebrate Ato members could catalyze a reaction similar to carbamoyl-AMP synthesis. However, its exact biochemical function in vertebrates needs further investigation. Another hypothesis is that a fusion of an enzyme with a functionally related transporter may directly link substrate import to its metabolic assimilation, enhancing the rate and/or efficiency of downstream enzymatic reactions.

In summary, this work provides new insights into the structure and potential function of Ato members in *C. albicans* but also serves as a basis for further studies in higher eukaryotes.

## 5. Funding & Acknowledgements

This work was supported by the MetaFungal project PTDC/BIA-MIC/5246/2020 (http://doi.org/10.54499/PTDC/BIA-MIC/5246/2020) through the ‘Fundação para a Ciência e a Tecnologia’ (FCT).

Work at University of Minho (CBMA) was supported by the ‘Contrato-Programa’ UID/04050 funded by national funds through the FCT I.P.

Work at KU Leuven (Laboratory of Molecular Cell Biology) was supported by a grant from the Fund of Scientific Research Flanders (FWO # G0C0622N).

FG acknowledges FCT for the 2023.03135.BD PhD grant (https://doi.org/10.54499/2023.03135.BD) and Erasmus+ Program for supporting her stay at KU Leuven (Belgium).

PA acknowledges FCT for the 2024.03178.BDANA PhD grant.

JA acknowledges FCT for the PD/BD/150584/2020 (https://doi.org/10.54499/PD/BD/150584/2020).

AG-G acknowledges FCT for the 2021.08564.BD PhD grant (https://doi.org/10.54499/2021.08564.BD). WVG was supported by the KU Leuven Research Council grant (# C14/22/075).

AJPB was supported by a programme grant from the UK Medical Research Council [MR/M026663/2], a Wellcome Investigator Award (224323/Z/21/Z), the Medical Research Council Centre for Medical Mycology at the University of Exeter (MR/N006364/2 and MR/V033417/1), and the NIHR Exeter Biomedical Research Centre.

## 7. Supporting Information

This article contains supporting information files:

- Supplementary Data_1 (PDF file)

## 8. Author Contributions

**FG** performed the lab experiments and was responsible for the *in-silico* analysis, curation, analysis and visualisation of the data. **CBA** was actively involved in this work, and made important contributions to the experiments, particularly in Western blot, analysis and visualisation of data. **YP** performed Bioinformatics and phylogenetic analysis. **PA** was actively involved in the lab work and made important contributions to the experiments, particularly in Western blot and DNA cloning. **JA, VF** and **MC** contributed to the Bioinformatics analysis. **RA** constructed some of the mutant strains. **AG, WVG, JN** and **PVD** contributed to construction of GFP fusions. **ISS**, **AAP**, **GD** and **AJPB** contributed to conceptualization, validation, and supervision. **SP** and **PVD** contributed to funding acquisition, conceptualization, validation and supervision. **FG** drafted the original manuscript with additional input from all authors.

## References

1. Parambath S, Dao A, Kim HY, Zawahir S, Alastruey Izquierdo A, Tacconelli E, et al. *Candida albicans*— A systematic review to inform the World Health Organization Fungal Priority Pathogens List. Medical Mycology. 2024 Jun 1;62(6):myae045.

2. Katsipoulaki M, Stappers MHT, Malavia-Jones D, Brunke S, Hube B, Gow NAR. *Candida albicans* and *Candida glabrata*: global priority pathogens. Microbiology and Molecular Biology Reviews [Internet]. 2024 Jun 27 [cited 2025 Apr 15]; Available from: https://journals.asm.org/doi/10.1128/mmbr.00021-23

3. Lass-Flörl C, Steixner S. The changing epidemiology of fungal infections. Molecular Aspects of Medicine. 2023 Dec 1;94:101215.

4. Osei Sekyere J. *Candida auris*: A systematic review and meta-analysis of current updates on an emerging multidrug-resistant pathogen. MicrobiologyOpen. 2018;7(4):e00578.

5. Diekema D, Arbefeville S, Boyken L, Kroeger J, Pfaller M. The changing epidemiology of healthcare-associated candidemia over three decades. Diagnostic Microbiology and Infectious Disease. 2012 May 1;73(1):45–8.

6. Pfaller MA, Diekema DJ. Epidemiology of Invasive Candidiasis: a Persistent Public Health Problem. Clinical Microbiology Reviews. 2007 Jan;20(1):133–63.

7. Singh S, Sobel JD, Bhargava P, Boikov D, Vazquez JA. Vaginitis Due to *Candida krusei:* Epidemiology, Clinical Aspects, and Therapy. Clinical Infectious Diseases. 2002 Nov 1;35(9):1066–70.

8. Alves R, Sousa-Silva M, Vieira D, Soares P, Chebaro Y, Lorenz MC, et al. Carboxylic acid transporters in *candida* pathogenesis. mBio. 2020;11(3):1–11.

9. Pappas PG, Lionakis MS, Arendrup MC, Ostrosky-Zeichner L, Kullberg BJ. Invasive candidiasis. Nat Rev Dis Primers. 2018 May 11;4(1):1–20.

10. Tsai MH, Hsu JF, Yang LY, Pan YB, Lai MY, Chu SM, et al. Candidemia due to uncommon *Candida* species in children: new threat and impacts on outcomes. Sci Rep. 2018 Oct 15;8(1):15239.

11. Bougnoux ME, Brun S, Zahar JR. Healthcare-associated fungal outbreaks: New and uncommon species, New molecular tools for investigation and prevention. Antimicrob Resist Infect Control. 2018 Mar 27;7(1):45.

12. Kohlenberg A, Struelens MJ, Monnet DL, Plachouras D. *Candida auris:* epidemiological situation, laboratory capacity and preparedness in European Union and European Economic Area countries, 2013 to 2017. Euro Surveill. 2018 Mar 29;23(13):18–00136.

13. Alves R, Barata-Antunes C, Casal M, Brown AJP, Dijck PV, Paiva S. Adapting to survive: How *Candida* overcomes host-imposed constraints during human colonization. PLOS Pathogens. 2020 May 21;16(5):e1008478.

14. Cauchie M, Desmet S, Lagrou K. *Candida* and its dual lifestyle as a commensal and a pathogen. Research in Microbiology. 2017 Nov 1;168(9):802–10.

15. Mayer FL, Wilson, Duncan, and Hube B. *Candida albicans* pathogenicity mechanisms. Virulence. 2013 Feb 15;4(2):119–28.

16. Ene IV, Brunke S, Brown AJP, Hube B. Metabolism in Fungal Pathogenesis. Cold Spring Harb Perspect Med. 2014 Dec 1;4(12):a019695.

17. Childers DS, Raziunaite I, Avelar GM, Mackie J, Budge S, Stead D, et al. The Rewiring of Ubiquitination Targets in a Pathogenic Yeast Promotes Metabolic Flexibility, Host Colonization and Virulence. PLOS Pathogens. 2016 Apr 13;12(4):e1005566.

18. Johnston M. Feasting, fasting and fermenting: glucose sensing in yeast and other cells. Trends in Genetics. 1999 Jan 1;15(1):29–33.

19. impact of the Fungus-Host-Microbiota interplay upon *Candida albicans* infections: current knowledge and new perspectives | FEMS Microbiology Reviews | Oxford Academic [Internet]. [cited 2025 Apr 15]. Available from: https://academic.oup.com/femsre/article/45/3/fuaa060/6000215

20. Padder SA, Ramzan, Asiya, Tahir, Inayatullah, Rehman, Reiaz ul, and Shah AH. Metabolic flexibility and extensive adaptability governing multiple drug resistance and enhanced virulence in *Candida albicans*. Critical Reviews in Microbiology. 2022 Jan 2;48(1):1–20.

21. Morrison DJ, and Preston T. Formation of short chain fatty acids by the gut microbiota and their impact on human metabolism. Gut Microbes. 2016 May 3;7(3):189–200.

22. The *Candida albicans* ATO Gene Family Promotes Neutralization of the Macrophage Phagolysosome | Infection and Immunity [Internet]. [cited 2025 Apr 15]. Available from: https://journals.asm.org/doi/full/10.1128/iai.00984-15

23. The carboxylic acid transporters Jen1 and Jen2 affect the architecture and fluconazole susceptibility of *Candida albicans* biofilm in the presence of lactate: Biofouling: Vol 33, No 10 [Internet]. [cited 2025 Apr 15]. Available from: https://www.tandfonline.com/doi/abs/10.1080/08927014.2017.1392514?casa_token=H3PeX9KfO5QAAAAA:HjLWFZk1zvSSt_a0NSv5VrLGZj4d0bgz_ft7SEGjexDDG68Y96ZpWZgmOrZGRDhLqFJBV4Nnt4IZ

24. 24. Transport of carboxylic acids in yeasts | FEMS Microbiology Reviews | Oxford Academic [Internet]. [cited 2025 Apr 15]. Available from: https://academic.oup.com/femsre/article/32/6/974/2683394

25. Casal M, Queirós O, Talaia G, Ribas D, Paiva S. Carboxylic Acids Plasma Membrane Transporters in Saccharomyces cerevisiae. In: Ramos J, Sychrová H, Kschischo M, editors. Yeast Membrane Transport [Internet]. Cham: Springer International Publishing; 2016 [cited 2025 Apr 15]. p. 229–51. Available from: 10.1007/978-3-319-25304-6_9

26. Ribas D, Soares-Silva I, Vieira D, Sousa-Silva M, Sá-Pessoa J, Azevedo-Silva J, et al. The acetate uptake transporter family motif “NPAPLGL(M/S)” is essential for substrate uptake. Fungal Genetics and Biology. 2019 Jan 1;122:1–10.

27. Rabitsch KP, Tóth A, Gálová M, Schleiffer A, Schaffner G, Aigner E, et al. A screen for genes required for meiosis and spore formation based on whole-genome expression. Current Biology. 2001 Jul 10;11(13):1001–9.

28. Sá-Pessoa J, Amillis S, Casal M, Diallinas G. Expression and specificity profile of the major acetate transporter AcpA in *Aspergillus nidulans*. Fungal Genetics and Biology. 2015 Mar 1;76:93–103.

29. Tzschoppe K, Augstein A, Bauer R, Kohlwein SD, Barth G. trans-dominant mutations in the GPR1 gene cause high sensitivity to acetic acid and ethanol in the yeast *Yarrowia lipolytica*. Yeast. 1999 Nov 1;15(15):1645–56.

30. Paiva S, Devaux F, Barbosa S, Jacq C, Casal M. Ady2p is essential for the acetate permease activity in the yeast *Saccharomyces cerevisiae*. Yeast. 2004 Feb 1;21(3):201–10.

31. Sá-Pessoa J, Paiva S, Ribas D, Silva IJ, Viegas SC, Arraiano CM, et al. SATP (YaaH), a succinate–acetate transporter protein in *Escherichia coli*. Biochemical Journal. 2013 Aug 29;454(3):585–95.

32. Rendulić T, Alves J, Azevedo-Silva J, Soares-Silva I, Casal M. New insights into the acetate uptake transporter (AceTr) family: Unveiling amino acid residues critical for specificity and activity. Computational and Structural Biotechnology Journal. 2021;19:4412–25.

33. Sun P, Li J, Zhang X, Guan Z, Xiao Q, Zhao C, et al. Crystal structure of the bacterial acetate transporter SatP reveals that it forms a hexameric channel. Journal of Biological Chemistry. 2018;293(50):19492– 500.

34. Qiu B, Xia B, Zhou Q, Lu Y, He M, Hasegawa K, et al. Succinate-acetate permease from *Citrobacter koseri* is an anion channel that unidirectionally translocates acetate. Cell Research. 2018;28(6):644– 54.

35. Genomic Expression Programs in the Response of Yeast Cells to Environmental Changes | Molecular Biology of the Cell [Internet]. [cited 2025 Apr 15]. Available from: https://www.molbiolcell.org/doi/full/10.1091/mbc.11.12.4241

36. Boer VM, Winde JH de, Pronk JT, Piper MDW. The Genome-wide Transcriptional Responses of *Saccharomyces cerevisiae* Grown on Glucose in Aerobic Chemostat Cultures Limited for Carbon, Nitrogen, Phosphorus, or Sulfur *. Journal of Biological Chemistry. 2003 Jan 31;278(5):3265–74.

37. Danhof HA LM. The *Candida albicans* ATO Gene Family Promotes Neutralization of the Macrophage Phagolysosome. Infect Immun. 2015;83:4416–26.

38. Slavena V, J. CA, A. DH, R. CJ, Huaijin Z, C. LM. The Fungal Pathogen *Candida albicans* Autoinduces Hyphal Morphogenesis by Raising Extracellular pH. mBio. 2011 May 17;2(3):e00055–11.

39. MAFFT: a novel method for rapid multiple sequence alignment based on fast Fourier transform | Nucleic Acids Research | Oxford Academic [Internet]. [cited 2025 Apr 22]. Available from: https://academic.oup.com/nar/article/30/14/3059/2904316

40. Capella-Gutiérrez S, Silla-Martínez JM, Gabaldón T. trimAl: a tool for automated alignment trimming in large-scale phylogenetic analyses. Bioinformatics. 2009 Aug 1;25(15):1972–3.

41. IQ-TREE 2: New Models and Efficient Methods for Phylogenetic Inference in the Genomic Era | Molecular Biology and Evolution | Oxford Academic [Internet]. [cited 2025 Apr 22]. Available from: https://academic.oup.com/mbe/article/37/5/1530/5721363

42. Huerta-Cepas J, Serra F, Bork P. ETE 3: Reconstruction, Analysis, and Visualization of Phylogenomic Data. Molecular Biology and Evolution. 2016 Jun 1;33(6):1635–8.

43. Sievers F, Wilm A, Dineen D, Gibson TJ, Karplus K, Li W, et al. Fast, scalable generation of high-quality protein multiple sequence alignments using Clustal Omega. Mol Syst Biol. 2011 Oct 11;7:539.

44. Jumper J, Evans R, Pritzel A, Green T, Figurnov M, Ronneberger O, et al. Highly accurate protein structure prediction with AlphaFold. Nature. 2021;596(7873):583–9.

45. Varadi M, Anyango S, Deshpande M, Nair S, Natassia C, Yordanova G, et al. AlphaFold Protein Structure Database: Massively expanding the structural coverage of protein-sequence space with high-accuracy models. Nucleic Acids Research. 2022;50(D1):D439–44.

46. Smart OS, Neduvelil JG, Wang X, Wallace BA, Sansom MS. HOLE: a program for the analysis of the pore dimensions of ion channel structural models. J Mol Graph. 1996 Dec;14(6):354–60, 376.

47. Humphrey W, Dalke A, Schulten K. VMD: visual molecular dynamics. J Mol Graph. 1996 Feb;14(1):33–8, 27–8.

48. Wu M, Sun L, Zhou Q, Peng Y, Liu Z, Zhao S. Molecular Mechanism of Acetate Transport through the Acetate Channel SatP. Journal of Chemical Information and Modeling. 2019;59(5):2374–82.

49. Trott O, Olson AJ. AutoDock Vina: Improving the speed and accuracy of docking with a new scoring function, efficient optimization, and multithreading. Journal of Computational Chemistry. 2010 Jan 30;31(2):455–61.

50. Ribas D, A-Pessoa JS, Soares-Silva I, Paiva S, Nygård Y, Nygård N, et al. Yeast as a tool to express sugar acid transporters with biotechnological interest. FEMS Yeast Research. 2017;17:5.

51. Blom N, Gammeltoft S, Brunak S. Sequence and structure-based prediction of eukaryotic protein phosphorylation sites1. Journal of Molecular Biology. 1999 Dec 17;294(5):1351–62.

52. Prediction of post-translational glycosylation and phosphorylation of proteins from the amino acid sequence - Blom - 2004 - PROTEOMICS - Wiley Online Library [Internet]. [cited 2025 May 24]. Available from: https://analyticalsciencejournals.onlinelibrary.wiley.com/doi/abs/10.1002/pmic.200300771

53. Nguyen N, Quail MMF, Hernday AD. An Efficient, Rapid, and Recyclable System for CRISPR-Mediated Genome Editing in *Candida albicans*. mSphere. 2017;2(2).

54. Van Genechten W, Demuyser L, Dedecker P, Van Dijck P. Presenting a codon-optimized palette of fluorescent proteins for use in *Candida albicans*. Scientific Reports. 2020;10(1):1–9.

55. Min K, Ichikawa Y, Woolford CA, Mitchell AP. *Candida albicans* Gene Deletion with a Transient CRISPR- Cas9 System. mSphere. 2016;1(3):e00130–16.

56. Walther A, Wendland J. An improved transformation protocol for the human fungal pathogen *Candida albicans*. Curr Genet. 2003 Mar;42(6):339–43.

57. Volland C, Urban-Grimal D, Géraud G, Haguenauer-Tsapis R. Endocytosis and degradation of the yeast uracil permease under adverse conditions. Journal of Biological Chemistry. 1994 Apr 1;269(13):9833– 41.

58. Sousa-Silva M, Soares P, Alves J, Vieira D, Casal M, Soares-Silva I. Uncovering Novel Plasma Membrane Carboxylate Transporters in the Yeast *Cyberlindnera jadinii*. Journal of Fungi. 2022;8(1).

59. Baldi N, De Valk SC, Sousa-Silva M, Casal M, Soares-Silva I, Mans R. Evolutionary engineering reveals amino acid substitutions in Ato2 and Ato3 that allow improved growth of *Saccharomyces cerevisiae* on lactic acid. FEMS Yeast Research. 2021;21(4):1–12.

60. Reinders J, Wagner K, Zahedi RP, Stojanovski D, Eyrich B, van der Laan M, et al. Profiling Phosphoproteins of Yeast Mitochondria Reveals a Role of Phosphorylation in Assembly of the ATP Synthase *. Molecular & Cellular Proteomics. 2007 Nov 1;6(11):1896–906.

61. Řičicová M, Kučerová H, Váchová L, Palková Z. Association of putative ammonium exporters Ato with detergent-resistant compartments of plasma membrane during yeast colony development: pH affects Ato1p localisation in patches. Biochimica et Biophysica Acta (BBA) - Biomembranes. 2007 May 1;1768(5):1170–8.

62. Strachotová D, Holoubek A, Kučerová H, Benda A, Humpolíčková J, Váchová L, et al. Ato protein interactions in yeast plasma membrane revealed by fluorescence lifetime imaging (FLIM). Biochimica et Biophysica Acta (BBA) - Biomembranes. 2012 Sep 1;1818(9):2126–34.

63. Thiaville PC, El Yacoubi B, Köhrer C, Thiaville JJ, Deutsch C, Iwata-Reuyl D, et al. Essentiality of threonylcarbamoyladenosine (t6A), a universal tRNA modification, in bacteria. Molecular Microbiology. 2015;98(6):1199–221.

64. Murphy FV, Ramakrishnan V, Malkiewicz A, Agris PF. The role of modifications in codon discrimination by tRNALysUUU. Nat Struct Mol Biol. 2004 Dec;11(12):1186–91.

65. Lin CA, Ellis, Steven R., and True HL. The Sua5 Protein Is Essential for Normal Translational Regulation in Yeast. Molecular and Cellular Biology. 2010 Jan;30(1):354–63.

