## Supporting information 1 for "Expansion, functional diversification and gene fusion events in the Ato protein family"

Tables S1

Figures S1, S2, S3, S4, S5, S6, S7, S8, S9

**Table S1. List of oligonucleotides used in this work.**

| Name | Sequence |
|---|---|
| Hernday-AHO1096 | GACGGCACGGCCACGCGTTTAAACCGCC |
| Hernday-AHO1098 | CAAATTAAAAATAGTTTACGCAAG |
| Hernday-AHO1097 | CCCGCCAGGCGCTGGGGTTTAAACACCG |
| Hernday-AHO1237 | AGGTGATGCTGAAGCTATTGAAG |
| Hernday-AHO1236 | TAAAGCTGCCACAAGAGGTATTTC |
| Hernday-gRNA-ATO1-GFP | CGTAAACTATTTTTAATTTGGGATCTTGTAAATCTGGTAAGTTTTAGAGCTAGAAATAG |
| Hernday-gRNA-ATO2-GFP | CGTAAACTATTTTTAATTTGTCTGAAGTAGGAGTTTGTTGGTTTTAGAGCTAGAAATAG |
| Hernday-gRNA-ATO3-GFP | CGTAAACTATTTTTAATTTGGCAATGGAATAGAAACAGGTGTTTTAGAGCTAGAAATAG |
| Hernday-gRNA-ATO4-GFP | CGTAAACTATTTTTAATTTGTCTGGTAAAGGAATTTCTCTGTTTTAGAGCTAGAAATAG |
| Hernday-gRNA-ATO5-GFP | CGTAAACTATTTTTAATTTGTCTGGTAAGGGAATTTCTTTGTTTTAGAGCTAGAAATAG |
| Hernday-gRNA-ATO6-GFP | CGTAAACTATTTTTAATTTGCCAAATTGAGGGACTGGAAGGTTTTAGAGCTAGAAATAG |
| Hernday-gRNA-ATO7-GFP | CGTAAACTATTTTTAATTTGTTTTTGTTACCCCCAAGATGGTTTTAGAGCTAGAAATAG |
| Hernday-gRNA-ATO8-GFP | CGTAAACTATTTTTAATTTGTTTGCCTCACTTCCTGGCAAGTTTTAGAGCTAGAAATAG |
| Hernday-gRNA-ATO9-GFP | CGTAAACTATTTTTAATTTGGGGTAATTGTTGTATAATCGGTTTTAGAGCTAGAAATAG |
| Hernday-gRNA-ATO10-GFP | CGTAAACTATTTTTAATTTGCTTGGTTATCAAGACATCACGTTTTAGAGCTAGAAATAG |
| SNR52/F | AAGAAAGAAAGAAAACCAGGAGTGAA |
| SNR52/R_ATO1 | CCAAATGCTTCAACTAAATCCAAATTAAAAATAGTTTACGCAAGTC |
| SNR52/R_ATO2 | CCGAAAGCAGCCATCAATTCCAAATTAAAAATAGTTTACGCAAGTC |
| SNR52/R_ATO3 | CCGAAGGCAGCCATCAAGTCCAAATTAAAAATAGTTTACGCAAGTC |
| sgRNA/F_ATO1 | GATTTAGTTGAAGCATTTGGGTTTTAGAGCTAGAAATAGCAAGTTAAA |
| sgRNA/F_ATO2 | GAATTGATGGCTGCTTTCGGGTTTTAGAGCTAGAAATAGCAAGTTAAA |
| sgRNA/F_ATO3 | GACTTGATGGCTGCCTTCGGGTTTTAGAGCTAGAAATAGCAAGTTAAA |
| sgRNA/R | ACAAATATTTAAACTCGGGACCTGG |
| SNR52/N | GCGGCCGCAAGTGATTAGACT |
| sgRNA/N | GCAGCTCAGTGATTAAGAGTAAAGATGG |
| CaCas9/F | ATCTCATTAGATTTGGAACTTGTGGGTT |
| CaCas9/R | TTCGAGCGTCCCAAAACCTTCT |
| Hernday-dDNA-ATO1-GFP-FW | GGTATTGCTAATCCTCAAAATAGTTATATTACTGTTAAAGCTATTCCATTACCAGATTTACAAGATCCAACAAGAAAAAATAAAGGTG CTGGCGCAGGTGCTATGTCTAAAGGTGAAGAATT |
| Hernday-dDNA-ATO1-GFP-RV | TTGATTGATTGATTGATTGATTGATTGATTACTTGGCCAATTATTTTATTTGTACAATTCATCCA |
| Hernday-dDNA-ATO2-GFP-FW | TCACTGCTATCATCGCTTGGTATGTTGCTTTAGCAGGTACAGCCACTACAACAAACTCCTACTTCAGACCAATTTCCATTCCAATGCC AGGAAATGTTGCTTTTAAAAACGGTGCTGGCGCAGGTGCTATGTCTAAAGGTGAAGAATT |
| Hernday-dDNA-ATO2-GFP-RV | AGCATTAAAATTAGATAAGTAGCAACAACTGAAAGAGTTTTGCCATTATTTGTACAATTCATCCA |
| Hernday-dDNA-ATO3-GFP-FW | GTGGAATGCCTTAGCCGGTACTGCTACTCCAACCAACTCTTACTTTCAACCTGTTTCTATTCCATTGCCAGGTAACG TTGTTTTCAAGAAAGGTGCTGGCGCAGGTGCTATGTCTAAAGGTGAAGAATT |

| | |
|---|---|
| Hernday-dDNA-ATO3-GFP-RV | GATAGCCATGGTGAATGACAATAAAATGCAGCTAGACAGATATTTTTATTTGTACAATTCATCCA |
| Hernday-dDNA-ATO4-GFP-FW | GAAGCTTTTGCTGGGGTAGCTAATAGACAAAACTCTTACATGGTTCCCCGTGAGATACCTTTACCAGATCTCTCAGTCTGGCTTAAACGTAAAAAAGCTGTCAGCAAAACATCAAATGGTGCTGGCGCAGGTGCTATGTCTAAAGGTGAAGAATT |
| Hernday-dDNA-ATO4-GFP-RV | AATACTGAATCTTTAGGGTTATTGTAGCCGGGTTCCGTTTTTGAGTTATTTGTACAATTCATCCA |
| Hernday-dDNA-ATO5-GFP-FW | GAAGCTTTCGCTGGTGTGGCCAACACACATAATTCATATTTGGTTCCCAAGGAGATACCCTTACCAGATCTTTCACTTTGGTTTAAACGTAAAAAGTCGGAAAATCAACCATCAGAGGGTGCTGGCGCAGGTGCTATGTCTAAAGGTGAAGAATT |
| Hernday-dDNA-ATO5-GFP-RV | AAATGTTTCCAGAACAAGTTACAAACAAAAATATAACGTTATACTTTATTTGTACAATTCATCCA |
| Hernday-dDNA-ATO6-GFP-FW | ATTTCTGGTATGGCAGATAAATTCAATTCTTATTTCACAGTTCATCCATTACCTGTTCCTCAATTTGGAAAGAAAAATGGTGCTGGCGCAGGTGCTATGTCTAAAGGTGAAGAATT |
| Hernday-dDNA-ATO6-GFP-RV | CATGAATAATAGAATTCTAGGATGCACAGCATAGCTAGTTTAGATTTATTTGTACAATTCATCCA |
| Hernday-dDNA-ATO7-GFP-FW | GCTGATAGATATAATTCTTATTTTACTATCAATCCAGTTCCAATTCCACATCTTGGGGGTAACAAAAAACAAAAAGAGATGAGAGGTGCTGGCGCAGGTGCTATGTCTAAAGGTGAAGAATT |
| Hernday-dDNA-ATO7-GFP-RV | AGACAAGTTGACCTCCCCCTTTTTCTGTAGAACTCAGATCTCATCTTATTTGTACAATTCATCCA |
| Hernday-dDNA-ATO8-GFP-FW | TTATTGGCAACAAAACAGAACTCTTACTTTACACTTTCTGTGATTTCATTGCCAGGAAGTGAGGCAAAACAAAAGAAAGGTGCTGGCGCAGGTGCTATGTCTAAAGGTGAAGAATT |
| Hernday-dDNA-ATO8-GFP-RV | AATGAAAGAAATCCAAATAAACCGTTTTCCAGCTCTTAACCTTTGTTATTTGTACAATTCATCCA |
| Hernday-dDNA-ATO9-GFP-FW | TAGCTGGTGTTTGGGAATTTTTTGTGGGTAATTGTTGTATAATCGTTGCATTCGACTCAAATAAAGAACAGATTGCATTACCAATTGGTGCTGGCGCAGGTGCTATGTCTAAAGGTGAAGAATT |
| Hernday-dDNA-ATO9-GFP-RV | CTAATTTACGCTAAGGTGTTAGCTTATTGGACCAACTACTTTAAATTATTTGTACAATTCATCCA |
| Hernday-dDNA-ATO10-GFP-FW | AATTCATATTTTGTCATTCCTATCACTTGGTTATCAAGACATCACGGTAAGGGTGCTGGCGCAGGTGCTATGTCTAAAGGTGAAGAATT |
| Hernday-dDNA-ATO10-GFP-RV | AAGTTTGTTGTCTTCCCCCTTTGATTGTAGTTTTCCACTAACTTCTTATTTGTACAATTCATCCA |
| NAT_ATO1_repair/F | ACACTACAATAAACTTTTAACAACACACATAAATAACTAACTACTACAACTACAACTACAACCACTACACTTATCAAATCAGTCTAATCACTTGCGGCCGC |
| NAT_ATO1_repair/R | TCATCATTAAAAAAAAAAATCAATACCAGTATTGATTGATTGATTGATTGATTGATTGATTGATTACTTGGCCAATTATTGGACCACCTTTGATTGTAAATAG |
| NAT_ATO2_repair/F | CCAAATTGCTGTTCATTTGTTTTAGCTATTATTTTGTTCGTTCTACGAAGAATCTTATTATTGATCTACTGTTAAAGAATAGTCTAATCACTTGCGGCCGC |
| NAT_ATO2_repair/R | TTATAGTTGAATCGTACAATTGAACAATCAAATTAAGCATTAAAATTAGATAAGTAGCAACAACTGAAAGAGTTTTGCCAGGACCACCTTTGATTGTAAATAG |
| NAT_ATO3_repair/F | TTCGAAATAGAAAAAAACTGTTTCTTTTATATAAAGTTCTACTATCTATATCCAATATCGATAAATAAAAGGAAACAACTAGTCTAATCACTTGCGGCCGC |
| NAT_ATO3_repair/R | AATTATAGTAGCAATAAAGAAAAAAAAAATCAAATGATAGCCATGGTGAATGACAATAAAATGCAGCTAGACAGATATTTGGACCACCTTTGATTGTAAATAG |
| ATO1-FW-Check | ACAACTACAACTACAACCACTACACT |
| ATO1-RV-Check | ACACACACACACACACAGACCCA |
| ATO2-FW-Check | TCCAAGTTGGCCAAATTGCTGT |
| ATO2-RV-Check | GCAACAACTGAAAGAGTTTTGCCA |
| ATO3-FW-Check | TGATGTTTAGTTTTCACACCCCCA |
| ATO3-RV-Check | GCCATGGTGAATGACAATAAAATGCA |
| ATO4-FW-Check | GGCTATAAAAGGCCAACCAT |
| ATO4-RV-Check | TCGTCGTCAGGTAATTGCAG |
| ATO5-FW-Check | GGATGAACAGCCTTATTGTT |
| ATO5-RV-Check | GTTCGATCATCAACCTTCAC |
| ATO6-FW-Check | TTGTAGTTTCACTCAGGTTT |

| | |
|---|---|
| ATO6-RV-Check | TCCATGAGTTACCTCCACTG |
| ATO7-FW-Check | ACTCAGACCATCCATCCTCCTA |
| ATO7-RV-Check | ACCAGACAAGTTGACCTCCCCCT |
| ATO8-FW-Check | GTTACAATTGCAAACTGCTT |
| ATO8-RV-Check | AAGTAGTCGTGCATGTTTTC |
| ATO9-FW-Check | CCAATACCACTCTTTAATGT |
| ATO9-RV-Check | GGAAGGACGTGAAACATGAG |
| ATO10-FW-Check | CGTTCTAAATCTGTGTAGGC |
| ATO10-RV-Check | GCTAATTATGAGATAGCTTC |
| Inside-GFP-Check-RV | GTAATACCAGCAGCAGTAAC |

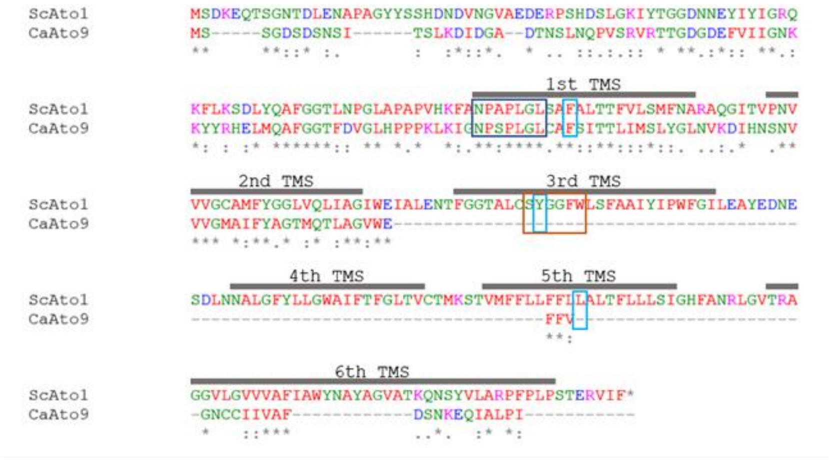


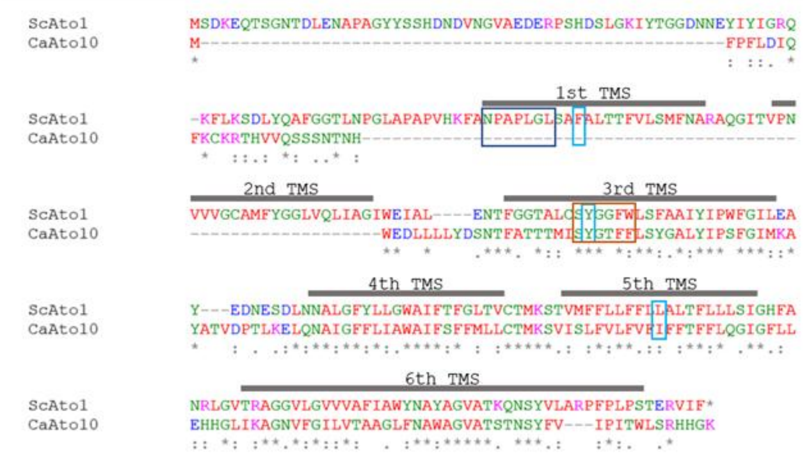


**Figure S1. Alignment of Ato9 and Ato10 protein sequences of *C. albicans* with Ato1 of *S. cerevisiae*.** The ClustalOmega was used to obtain the sequence alignment. The transmembrane regions of Ato1 of *S. cerevisiae* are highlighted in gray. The signature motif, N-P-A-P-L-G-L of the AceTr family is highlighted with dark blue rectangle. The narrowest hydrophobic constriction region (FLY) is represented by light blue rectangles. A second signature motif S-Y[F]-G-F-W located in the third TMS is highlighted in orange.

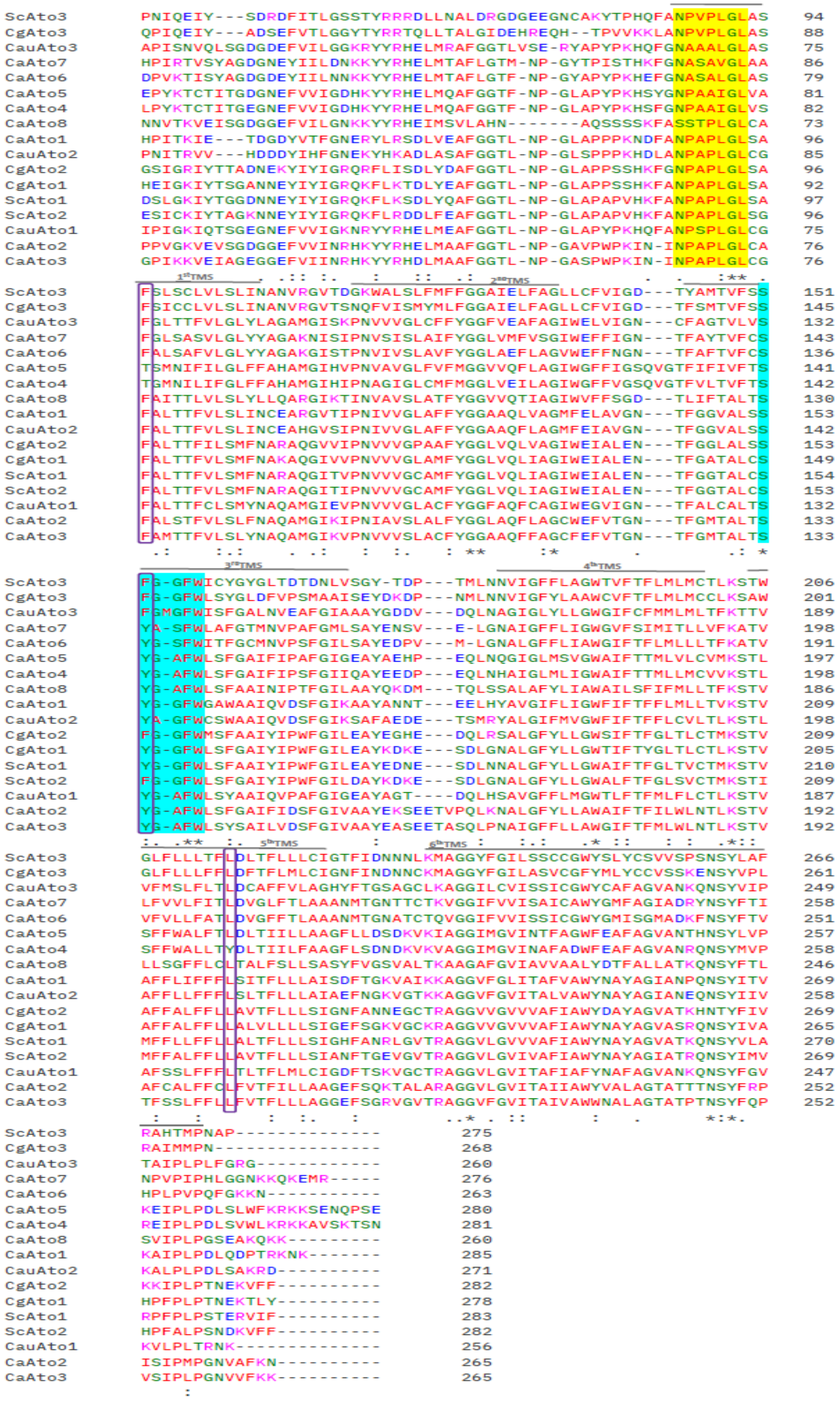


**Figure S2. Alignment of Ato homologous sequences from *C. albicans* (Ato1-8), *C. glabrata* (Ato1-3), *C. auris* (Ato1-3) and *S. cerevisiae* (Ato1-3).** The ClustalOmega was used to obtain the sequence alignment. The signature motif, N-P-A-P-L-G-L of the AceTr family is highlighted in yellow background. The narrowest hydrophobic constriction region (FLY) is represented by purple rectangles. A second signature motif S-Y[F]-G-F-W located in the third TMS is highlighted in blue.

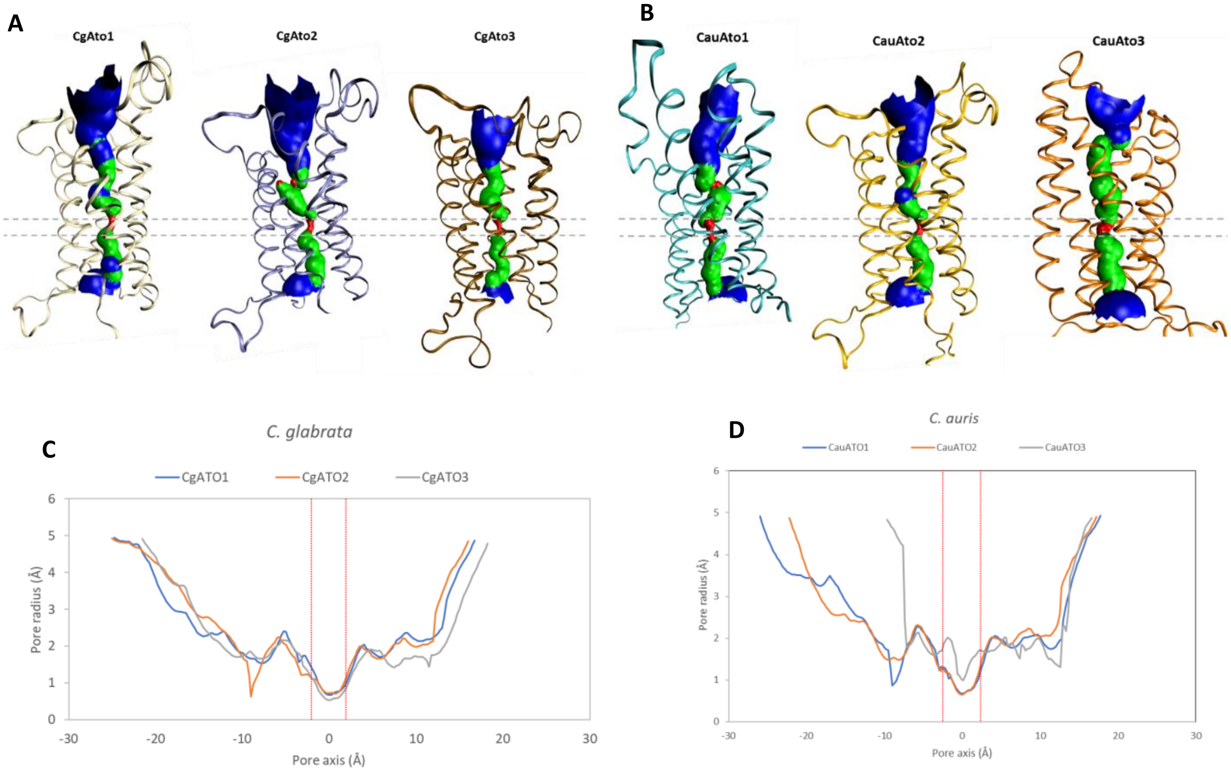


**Figure S3. Pore 3D structure predictions and radius profiles simulations along the channel axis in the Atos of *C. glabrate*, and *C. auris*. A.** Pore 3D structure prediction of CgAto1-3. **B.** Pore 3D structure prediction of CauAto1-3. The 3D structures of proteins are represented in New Ribbons, with the colour scheme consists of blue (representing a bigger pore size), green (representing an intermediate pore size), and red (representing a more constricted pore size) in pore prediction. The horizontal dashed grey lines correspond to the constricted site where the pore radius is tight. **C.** Simulations for the pore radius profiles along the channel axis in CgAto1-3. **D.** Simulations for the pore radius profiles along the channel axis in CauAto1-3. The region in the middle of proteins that contains the construction site is marked by vertical red dashed lines.

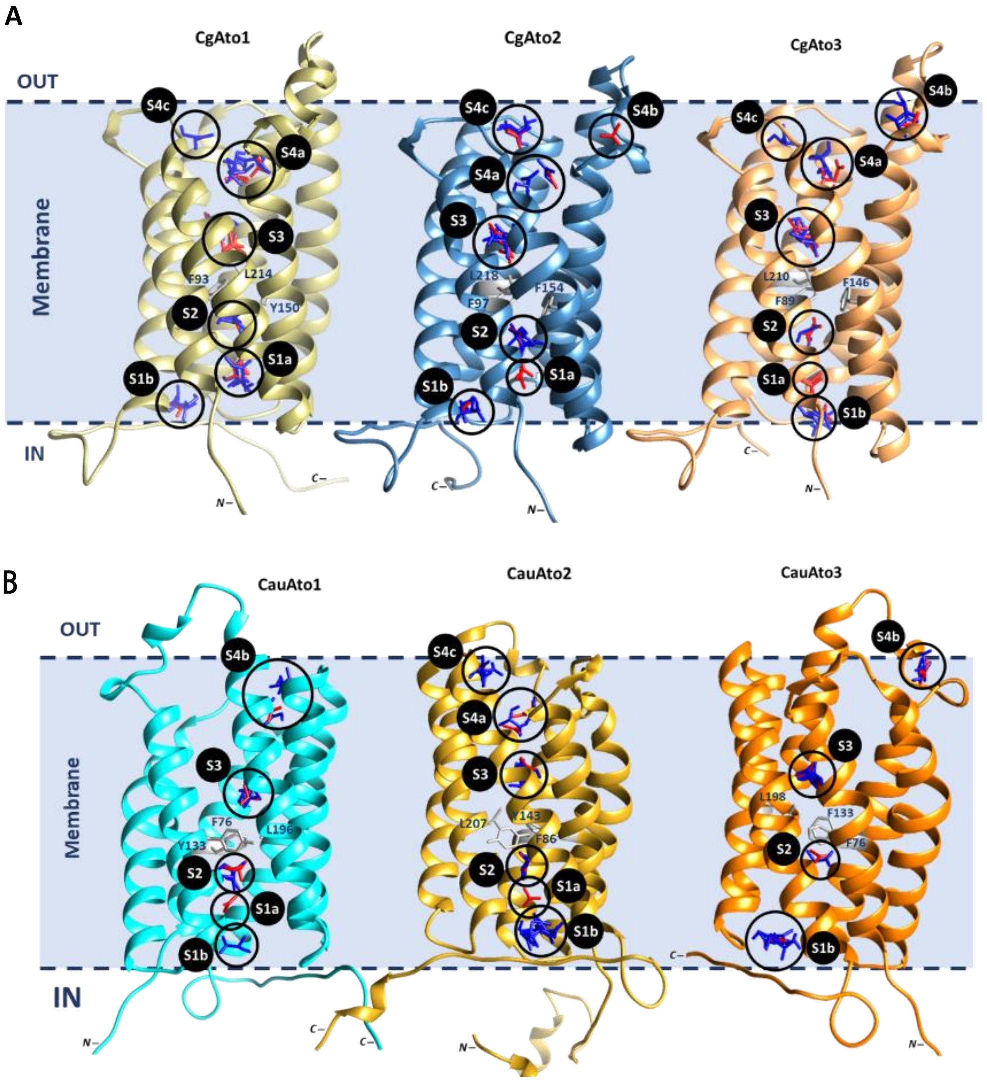


**Figure S4. 3D structure and molecular docking. A.** Molecular docking of CgAto1-3 with acetate and lactate as substrates. **B.** Molecular docking of CauAto1-3 with acetate and lactate as substrates. Predicted binding sites for acetate and lactate were shown with S1 to S4. Site S1 is located at the cytoplasmic vestibule, Sites S2 and S3 are located inside the main pore, and Site S4 is located at the periplasmic vestibule. Localization of the N- and C-terminal of the proteins is shown. Acetate and lactate ligands are presented in red and blue respectively.

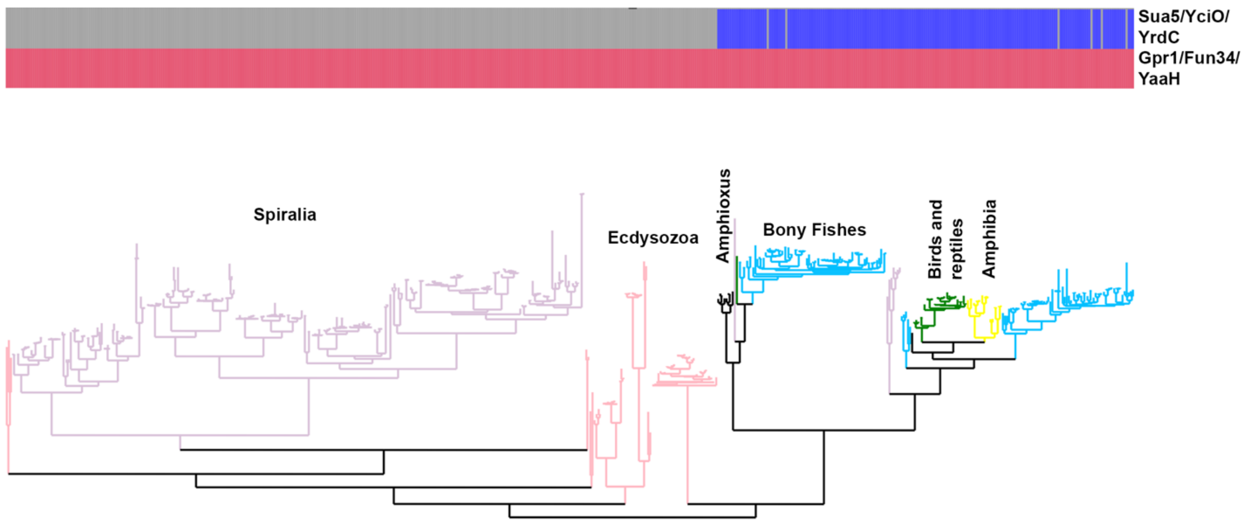


**Figure S5. Metazoan ATO phylogeny and structure.** Phylogeny and domain composition of metazoan ATO proteins. Domain presence is denoted as blue or red and domain absence with grey. Tree branch colors are colored based on taxonomy. Notice the presence of two clades: a protostomian and a mostly vertebrate clade.

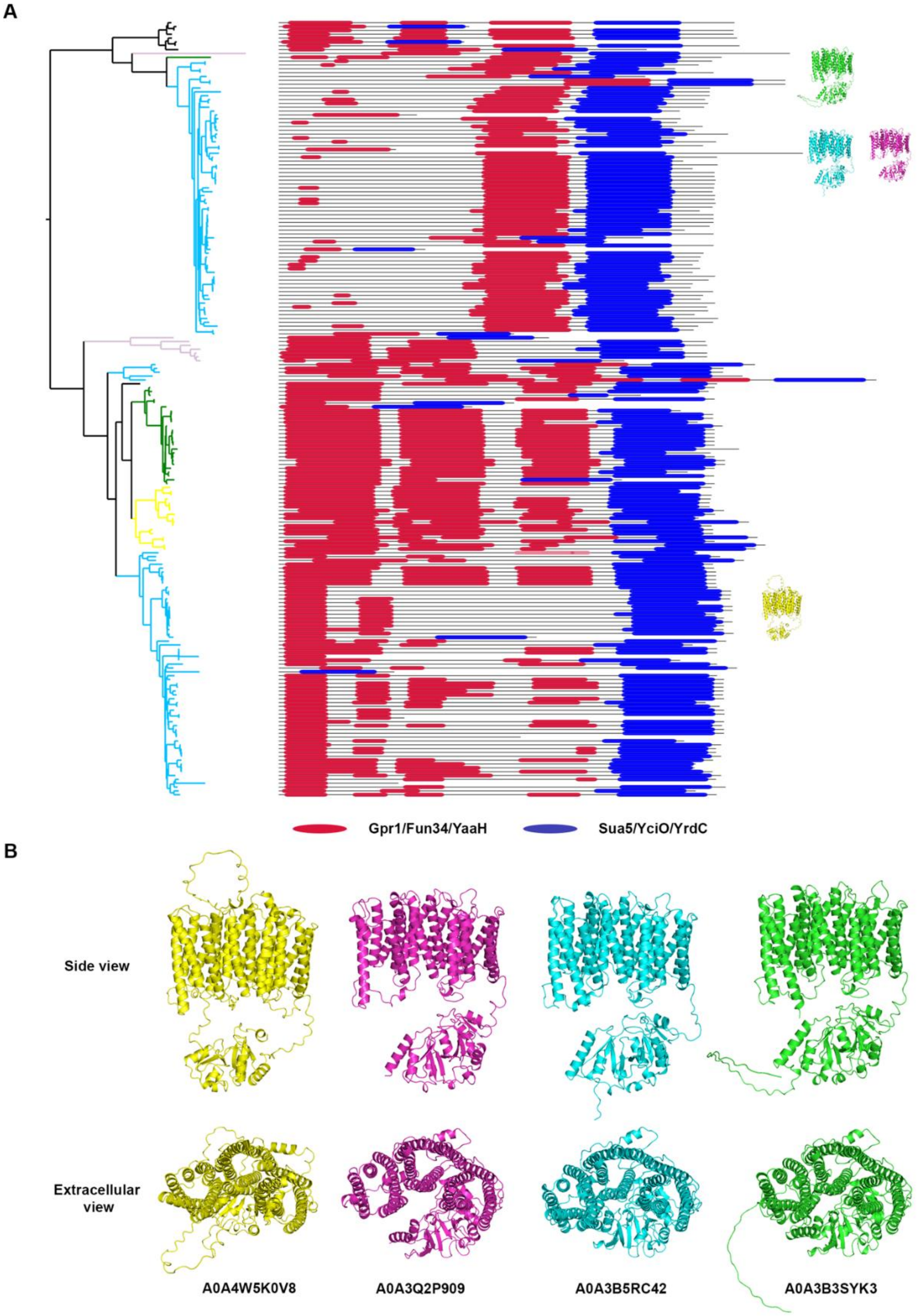


**Figure S6. Domain order of vertebrate ATOs and representative structures. A.** Domain order plotted next to the phylogenetic tree from Figure 6. Domains are plotted on a black line that is proportional to sequence length. Approximate phylogenetic position of the structures shown in B is presented next to the domain plot. **B.** Alphafold 2.0 structures of vertebrate ATO proteins (downloaded from Uniprot) that return less than three hits for the Gpr1/Fun34/YaaH domain. Notice that despite their divergent sequence the three copies of the characteristic six-helix SatP-fold are present in all of them.

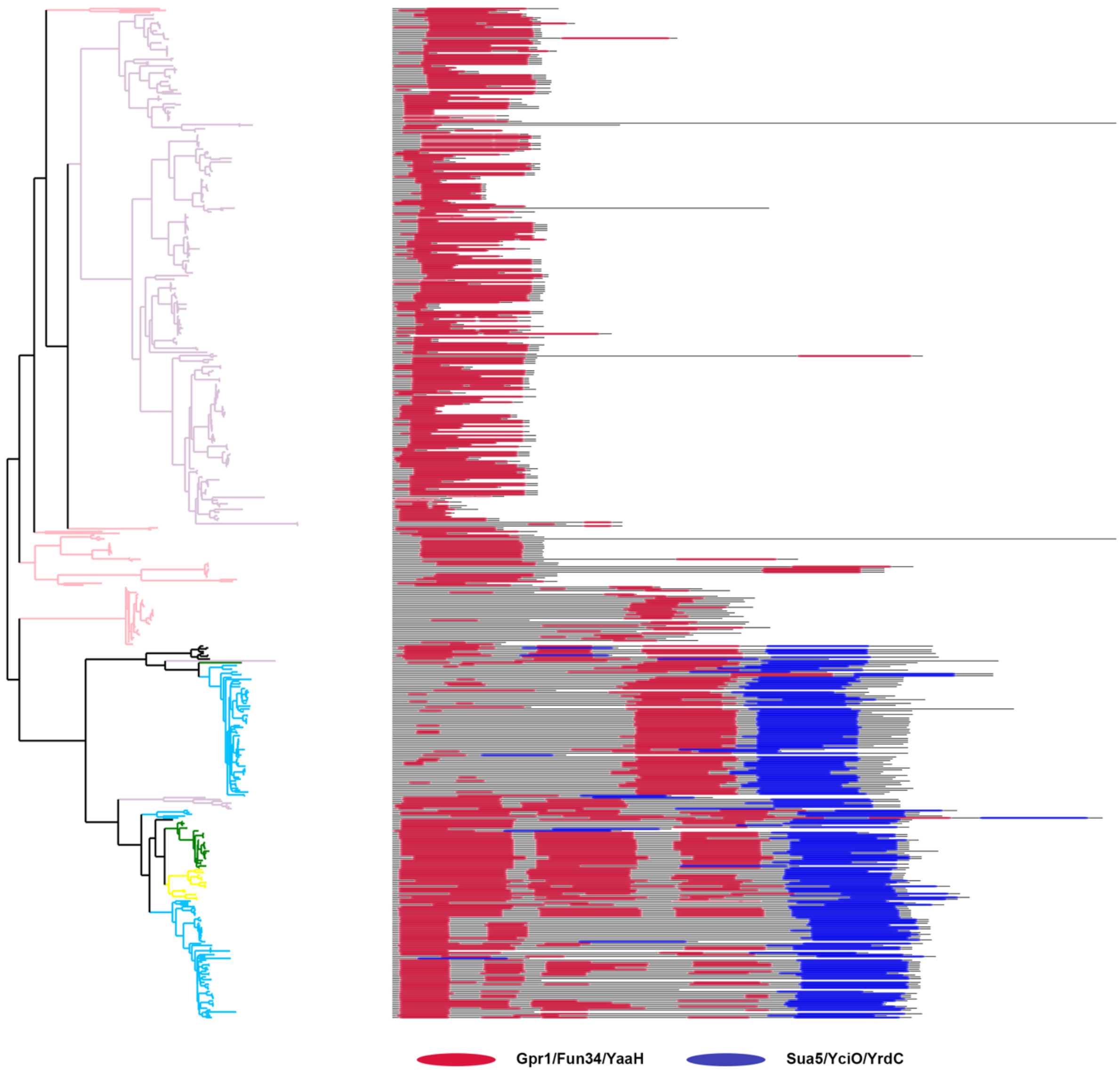


**Figure S7. Domain order of metazoan ATOs.** Domains are plotted as in Figure S6. Branch colors are the same as in figure S5. Notice that protostomian sequences are much shorter. They have only one copy of the Gpr1/Fun34/YaaH domain and lack the C-terminaly fused enzyme-like domain. Their structure is similar to their fungal and prokaryotic homologs.

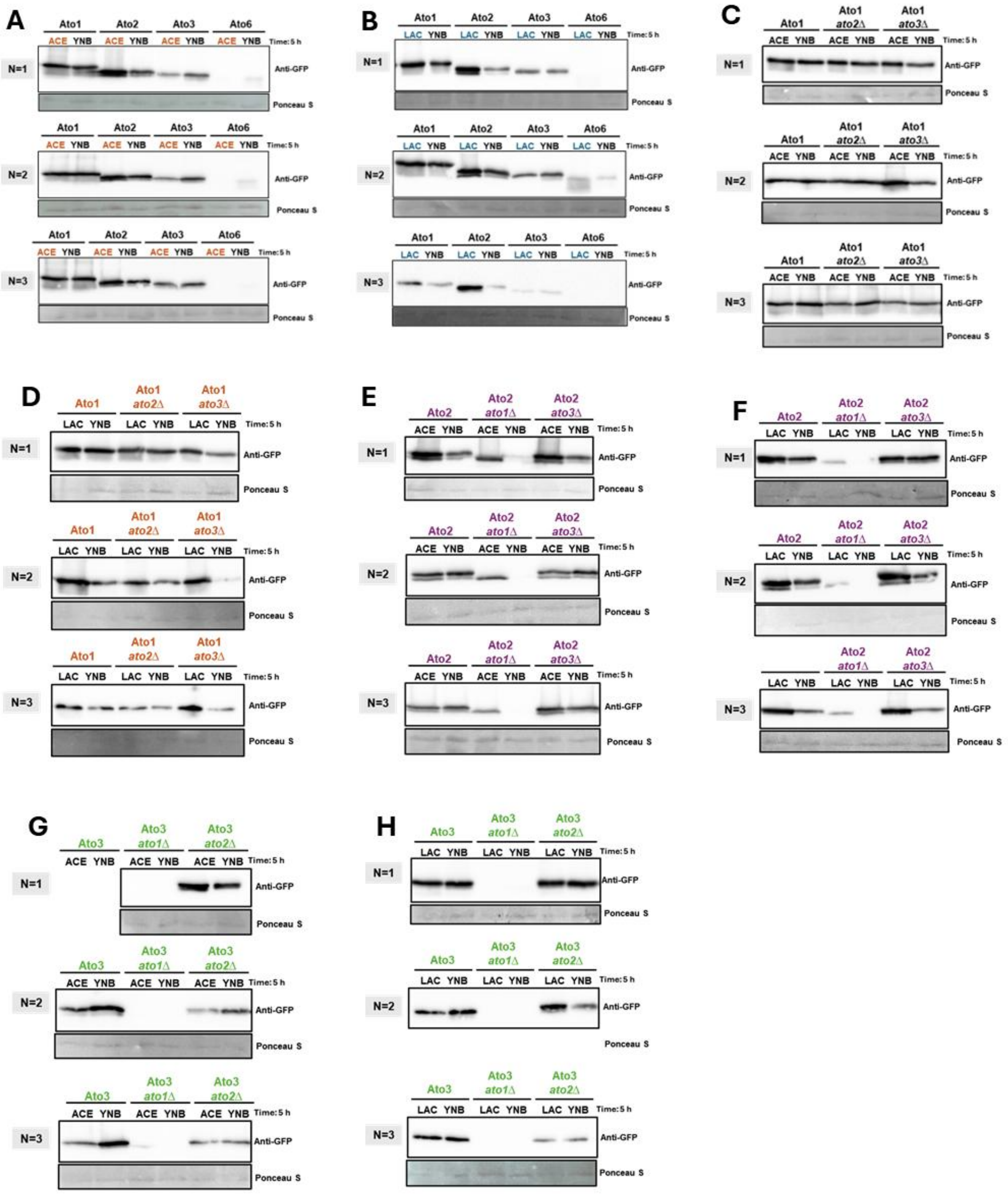


**Figure S8. Western Blots of the indicted Atos-GFP in *C. albicans* WT and *ato*-mutant cells.** *C. albicans* cells grown in 50 mL SM medium supplemented with 0.2 % (w/v) glucose to exponential phase. They were washed then with deionized water and then transferred, to fresh minimal media containing different carbon sources: 0.1 % (v/v) acetic acid pH 6.0 (ace), 0.1 % (v/v) lactic acid pH 5.0 (lac) and without any carbon source (YNB) (SM: 0.67% w/v YNB with ammonium sulphate). **A-H.** The cells were collected at the time-point 5h and total protein extracts were separated by SDS-PAGE. Atos-GFP proteins were detected with an anti-GFP antibody. Ponceau S staining was used as a loading control. The results represent three independent experiments, each using a different clone (n≥3).

```
# Sequence                  # x   Context     Score   Kinase    Answer
# -------------------------------------------------------------------
# Sequence                  3 S   --MPSTSSQ   0.534   PKC        YES
# Sequence                  4 T   -MPSTSSQK   0.753   PKC        YES
# Sequence                  6 S   PSTSSQKSV   0.921   unsp       YES
# Sequence                  9 S   SSQKSVGSS   0.996   unsp       YES
# Sequence                 13 S   SVGSSVMDP   0.988   unsp       YES
# Sequence                 28 S   KVEVSGDGG   0.958   unsp       YES
# Sequence                 54 T   AFGGTLNPG   0.507   cdc2       YES
# Sequence                 81 T   FALSTFVLS   0.678   PKC        YES
# Sequence                102 S   NIAVSLALF   0.616   PKA        YES
# Sequence                132 T   MTALTSYGA   0.835   unsp       YES
# Sequence                133 S   TALTSYGAF   0.837   PKC        YES
# Sequence                158 S   AYEKSEETV   0.571   CKI        YES
# Sequence                180 T   WAIFTFILW   0.647   PKC        YES
# Sequence                187 T   LWLNTLKST   0.833   unsp       YES
# Sequence                190 S   NTLKSTVAF   0.621   PKC        YES
# Sequence                214 S   AGEFSQKTA   0.744   unsp       YES
# Sequence                230 T   LGVITAIIA   0.502   PKG        YES
# Sequence                245 T   GTATTTNSY   0.593   PKC        YES
# Sequence                248 S   TTTNSYFRP   0.622   PKC        YES
# Sequence                254 S   FRPISIPMP   0.711   PKA        YES
#
    MPSTSSQKSVGSSVMDPNEPPVGKVEVSGDGGEFVVINRHKYYRHELMAA   #     50
    FGGTLNPGAVPWPKININPAPLGLCAFALSTFVLSLFNAQAMGIKIPNIA   #    100
    VSLALFYGGLAQFLAGCWEFVTGNTFGMTALTSYGAFWLSFGAIFIDSFG   #    150
    IVAAYEKSEETVPQLKNALGFYLLAWAIFTFILWLNTLKSTVAFCALFFC   #    200
    LFVTFILLAAGEFSQKTALARAGGVLGVITAIIAWYVALAGTATTTNSYF   #    250
    RPISIPMPGNVAFKN                                      #    300
%1  ..ST.S..S...S..............S......................   #     50
%1  ...T..........................T...................   #    100
%1  .S.............................TS.................   #    150
%1  .......S.....................T......T..S..........   #    200
%1  .............S...............T..............T..S..   #    250
%1  ...S............
```

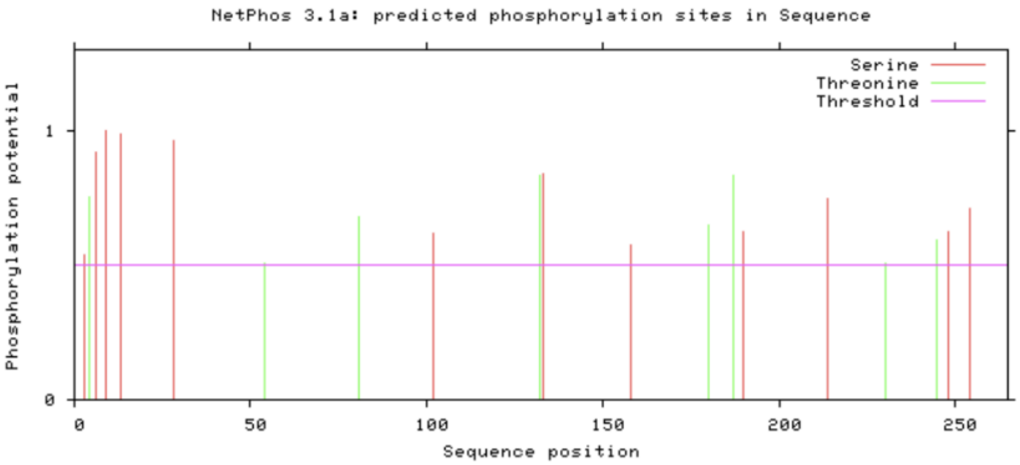


**Figure S9. Phosphorylation sites prediction in CaAto2 using Netphos-3.1b.** A total of 20 putative phosphorylation sites (prediction score ≥ 0.5) were identified, comprising 12 serine (indicated in red) and 8 threonine (indicated in green) residues. Four serine residues at positions 6, 9, 13, and 28 exhibited the highest prediction score (1.000). “#” indicates the position of the residue, and “X” represents the amino acid.
